# A population genetic analysis of the nematode *Strongyloides stercoralis* in Asia shows that human infection is not a zoonosis from dogs

**DOI:** 10.1101/2024.11.29.625877

**Authors:** Yuchen Liu, A. H. M. Raihan Sarker, Banchob Sripa, Sirikachorn Tangkawatta, Virak Khieu, William Nevin, Steve Paterson, Mark Viney

## Abstract

Gut nematode worms are important parasites of people and other animals. The parasitic nematode *Strongyloides stercoralis* infects an estimated 600 million people worldwide, and is one of the soil-transmitted helminthiases, a WHO-defined neglected tropical disease. It has long been suggested that human *S. stercoralis* infection may be a zoonosis from dogs. We investigated this by whole genome sequence analysis of *S. stercoralis* from sympatric human and dog populations in Asia. We find that human- and dog-derived *S. stercoralis* have genetically distinct nuclear genomes, but we also find evidence of rare cross infection. Analysis of the *S. stercoralis* mitochondrial genome reveals evidence of historical introgression between human- and dog-derived parasites. Based on these data we suggest that *S. stercoralis* was originally a parasite of canids, that began to infect humans when people domesticated dogs, since when human- and dog-derived parasites have differentiated, but have not become separate species.

**Significance Statement:** Nematode worms infect approximately a quarter of the world’s human population; *Strongyloides stercoralis* infects some 600 million people. It has been suggested that *S. stercoralis* in people and dogs is the same parasite, such that some human infection is acquired zoonotically from dogs. It is important to understand the source of human *Strongyloides* infection to be able to control it and the harm that it causes. Our population genomic analyses of human- and dog-derived *S. stercoralis* in Asia show that these parasites are distinct, but also reveal rare cross infection events. Our results are consistent with *S. stercoralis* ancestrally being a parasite of dogs that began infecting people when dogs became domesticated.

## Introduction

Parasitic nematodes are ubiquitous parasites of humans and other animals. In humans, soil-transmitted helminthiases (STH) are a collection of gut nematodes – *Ascaris*, hookworm, *Strongyloides* and *Trichuris* – that are transmitted through a common route where faeces from infected people contaminate the soil from which people acquire new infections by skin penetration of nematode larvae (*Strongyloides* and hookworm) or by ingesting parasite eggs (*Ascaris* and *Trichuris*). STHs infect over a billion people worldwide and are a World Health Organizaiton-defined neglected tropical disease (1).

*Strongyloides* is genus of *c.*60 species of parasitic nematode that infect a wide variety of mammals, birds, amphibians and reptiles (2). Two species of *Strongyloides* infect people, *S. stercoralis* and *S. fuelleborni. S. stercoralis* is widely distributed in tropical and sub-tropical regions, and is the most common species infecting people. *S. fuelleborni* was originally reported as a zoonosis from primates in Africa, but apparently zoonotic infection is now being reported in Asia (3, 4). *S. fuelleborni kellyi* is restricted to New Guinea and transmitted among people (in the absence of non-human primates from New Guinea) (5).

In the *Strongyloides* life cycle, hosts become infected by skin-penetrating larvae that develop into female only worms in the host gut. These reproduce parthenogenetically and their progeny leave the host in faeces (6). The progeny of a single parasitic female are genetically identical to each other and to their mother. Outside of the host the larvae can either develop directly into further infective larvae, or develop indirectly into free-living males and females that reproduce sexually (2, 7). *S. stercoralis* is unique among *Strongyloides* spp. in being able to internally autoinfect hosts (8).

It has long been hypothesised that *Strongyloides* in people and in dogs may be the same parasite, and thus that people may be zoonotically infected by dogs (9). This has been hypothesised because of the morphological similarity of *Strongyloides* from people and dogs, and because of case reports of people in non-*Strongyloides* endemic regions having acquired *S. stercoralis* infections, apparently from dogs (10, 11).

The question of a potential zoonotic origin of human *Strongyloides* infection is of practical importance. Specifically, if human infection includes a zoonotic origin then controlling human infection requires controlling the infection in both human and dogs. More generally, the question of the host range of parasitic nematodes is of basic biological interest, but little studied. Two other nematode parasites of people have zoonotic aspects to their transmission, *Ascaris* in pigs and *Dracunculus medinensis* in dogs. Population genetic analyses of *Ascaris* in pigs and people have found that they are an apparently interbreeding species complex but with distinct mitochondrial lineages (12,13). For *D. medinensis,* parasites in dogs and people are a single genetic population so that dogs are a zoonotic source of infection (14).

Previous studies have investigated whether *Strongyloides* in people and dog are the same parasites. Cross-infection experiments, where human-derived parasites have been used to infect dogs, have produced mixed results with some infections in dogs establishing, while others do not (9). Six studies have used a molecular approach, comparing *Strongyloides* from humans and dogs (4, 15–19). These have principally PCR amplified (either larvae isolated from faeces or faecal samples directly) fragments of the (i) nuclear 18s rRNA coding gene (focussing on two hypervariable regions) and (ii) mitochondrial *cox1* gene. These analyses have found (i) generally small levels (*e.g.* 4 variable positions in 11 nucleotides, and 2 in 11 nucleotides for two 18S hypervariable regions (16)) of genetic variation among samples, (ii) consistent evidence of differences between many *S. stercoralis* from people and dogs, but also (iii) evidence of *S. stercoralis* genotypes shared in parasites from people and dogs. A critique of these studies is that: (a) analyses of usually only single nuclear or mitochondrial loci, and detection of limited genetic variation limits the genetic resolution of *S. stercoralis* from people and dogs, (b) the within-species variability of the 18srRNA fragments (16), (c) that PCR amplification of faecal samples may lead to chimeric amplicons, and so the occurrence of these genotypes in individual larvae cannot be confirmed, and (d) that not all compare sympatric populations, and so geographical variation may confound the analyses (15).

Here we have used whole-genome, population genomics to investigate *S. stercoralis* infecting people and dogs. This builds on previous whole-genome population analyses of *S. ratti* in wild rats, which found that it consists of an assemblage of probably long-lived lineages, likely driven by its almost exclusive asexual mode of reproduction (20). The facultatively sexual reproduction in the *Strongyloides* life cycle can have significant effects on its population genetic structure, and can complicate making parasitological inferences from such data. In the present study we sampled *Strongyloides* from sympatric populations of people and dogs in Asia, whole genome sequenced individual larvae and used this to investigate the relationship between *S. stercoralis* infecting people and dogs, and its zoonotic potential.

## Results

### *S. stercoralis* occurs widely in people and dogs in Asia

We searched for *S. stercoralis* infection in sympatric communities of people and dogs in Bangladesh, Cambodia and Thailand (**Figure 1**). From screening 468 human and 394 dog faecal samples we detected an overall *S. stercoralis* prevalence of infection of 4.9% in people and 2.5% in dogs (**Table S1**). The prevalence differed among the countries, for people ranging from 1.4% in Bangladesh to 7.8% in Cambodia, and in dogs ranging from 0.7% in Thailand to 8.5% in Cambodia. These results are within the commonly reported ranges of *S. stercoralis* prevalence, though below the previous reports of high prevalence in Cambodia and Thailand (21, 22). In faeces *Strongyloides* can develop directly into infective third stage larvae (iL3s), or indirectly via free-living adults, whose progeny are iL3s. In Bangladesh we observed predominantly direct development; in Cambodia and Thailand human- and dog-derived faeces we observed free-living females, and so the iL3s we observed and collected likely developed by the indirect route.

**Figure 1.**
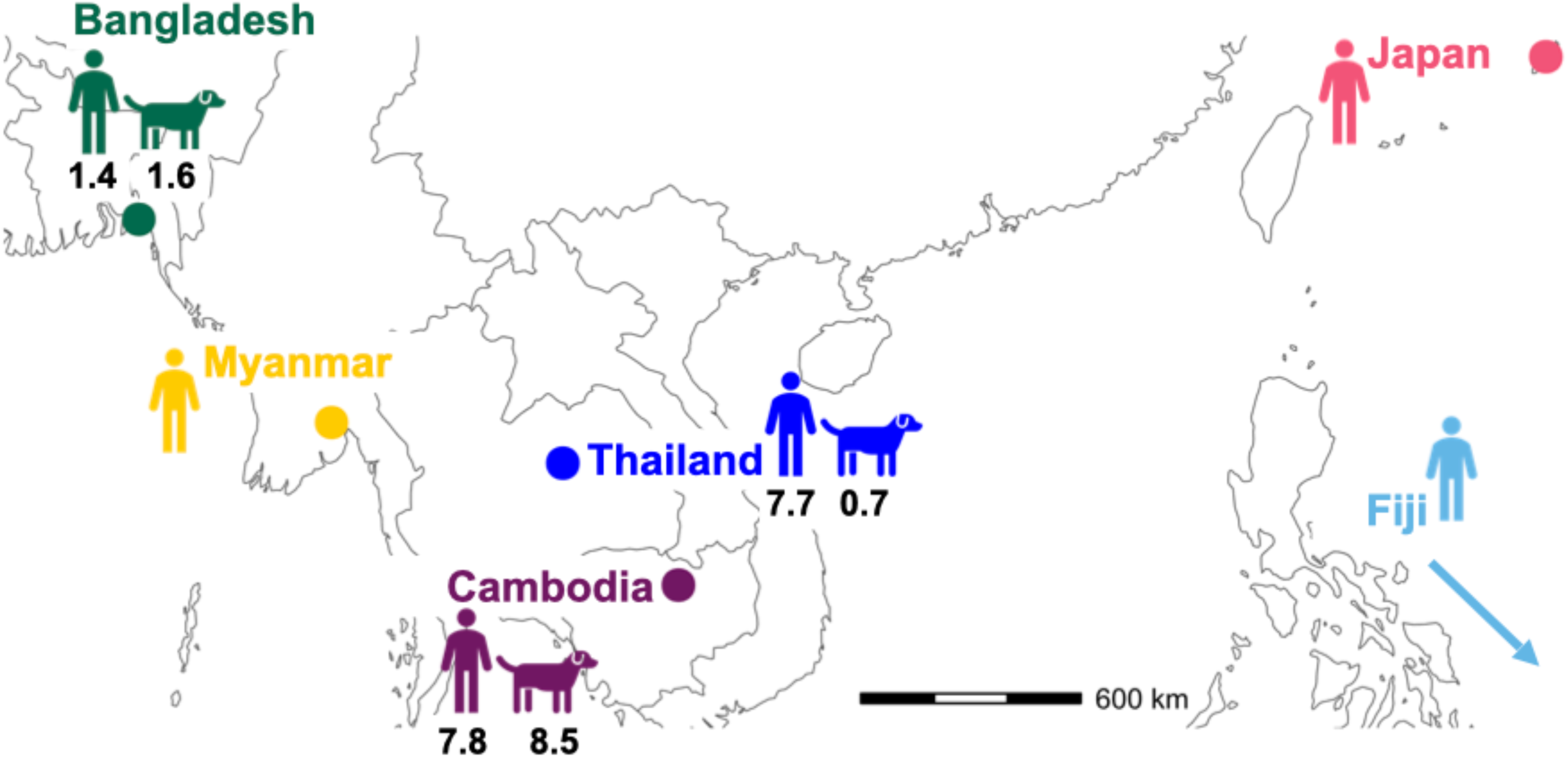
Distribution of sampling sites, and human and dog *S. stercoralis* prevalence. The prevalence of *S. stercoralis* in Bangladesh, Cambodia and Thailand in people and dogs, and the source of sequenced iL3s, including additional published data from Fiji, Myanmar and Japan.

From these Bangladesh, Cambodia and Thailand samples we collected and whole genome sequenced 207 iL3s from 26 people and 163 iL3s from 12 dogs, including 30 iL3s from three people who had migrated from Fiji to the UK (**Table S1**). A total of 276 iL3s (188 from people, 88 from dogs) produced sequence data that passed our quality control (QC) and which were identified as *S. stercoralis* by Kraken analysis. In a single dog faecal sample from Bangladesh 2 and 34 larvae were identified as a mixture of *S. ratti, S. venezuelensis*, respectively, (normally both parasites of rats) and *S. stercoralis* (**Table S1**). We presume that the non-*S. stercoralis* iL3s are contaminants when the human and dog samples were collected. To these 276 we added previously published whole genome amplification data for 12 iL3s from five people in Japan and 10 iL3s from three people in Myanmar (23), giving 298 samples in total. We aligned these sequence data to the *S. stercoralis* reference genome (PV001), which is an isolate of *S. stercoralis* derived from a dog in Pennsylvania, USA in the 1980s (24). A total of 143 alignments (109 from people, 34 from dogs) samples passed our quality control QC criteria of having an average read depth across the genome ≥ 20, and a ≥ 80% genome coverage, but the human and dog-derived samples differed in their alignment to the *S. stercoralis* reference. Specifically, for iL3s from people the mean depth and coverage was 46 (SD 23) and 98% (SD 0.04), respectively, but 29 (SD 8.7) and 88% (SD 0.04) for dog-derived iL3s (**Figure S1**).

A further 34 iL3s from seven people and 22 iL3s from four dogs produced sequence data that did not meet our QC criteria, and so we used a data combination strategy for closely related sequences from within individual hosts (**Figure S2; Table S2**). This resulted in an additional 10 human-derived samples and 5 dog-derived samples, giving a grand total of 158 samples, 119 from people and 39 from dogs (**Table S3**).

### *S. stercoralis* in people and dogs in Asia is genetically diverse

We identified 15,128 single nucleotide polymorphisms (SNPs) in the 158 samples (compared with the *S. stercoralis* 44.3 Mbp reference genome) giving an average SNP density of 3.4 SNPs / 10 kb, but with the density higher on the two autosomes (4.6 and 4.2 for chromones 1 and 2, respectively), compared with the X chromosome (1.7 across the six X chromosome scaffolds). For each of our samples, for each SNP we therefore knew what alleles were present (*i.e.* that there were 0, 1, or 2 copies of each variant), and we used these data to calculate the genetic distance between individual worm samples.

Considering all 158 samples, there was a positive association between genetic distance and geographical distance. This was greatest for the dog-derived samples from three countries (Mantel test R = 0.532, p = 0.001) compared with the human-derived samples from six countries (**Figure 1**), which covers a maximum geographical distance of *c.*10,400 km (Mantel test R = 0.299, p = 0.001).

### *S. stercoralis* from people and dogs have genetically distinct nuclear genomes

To investigate the relationship between human- and dog-derived *Strongyloides* we first analysed the fixation index, F_ST_. There was high F_ST_, thus high genetic differentiation, between human- and dog-derived iL3s (median F_ST_ = 0.43), as there was among the different dog-derived iL3s (median F_ST_ = 0.37), but much lower F_ST_ values among the human-derived iL3s (median F_ST_ = 0.16) (**Figure 2A; Table S4**). The 0.37 – 0.43 F_ST_ values are larger than would be expected from panmictically, sexually reproducing populations, which therefore suggests that these human- and dog-derived *Strongyloides* are genetically differentiated, as too are the dog-derived *Strongyloides* across Asia, and much more so than the human-derived *Strongyloides*.

**Figure 2.**
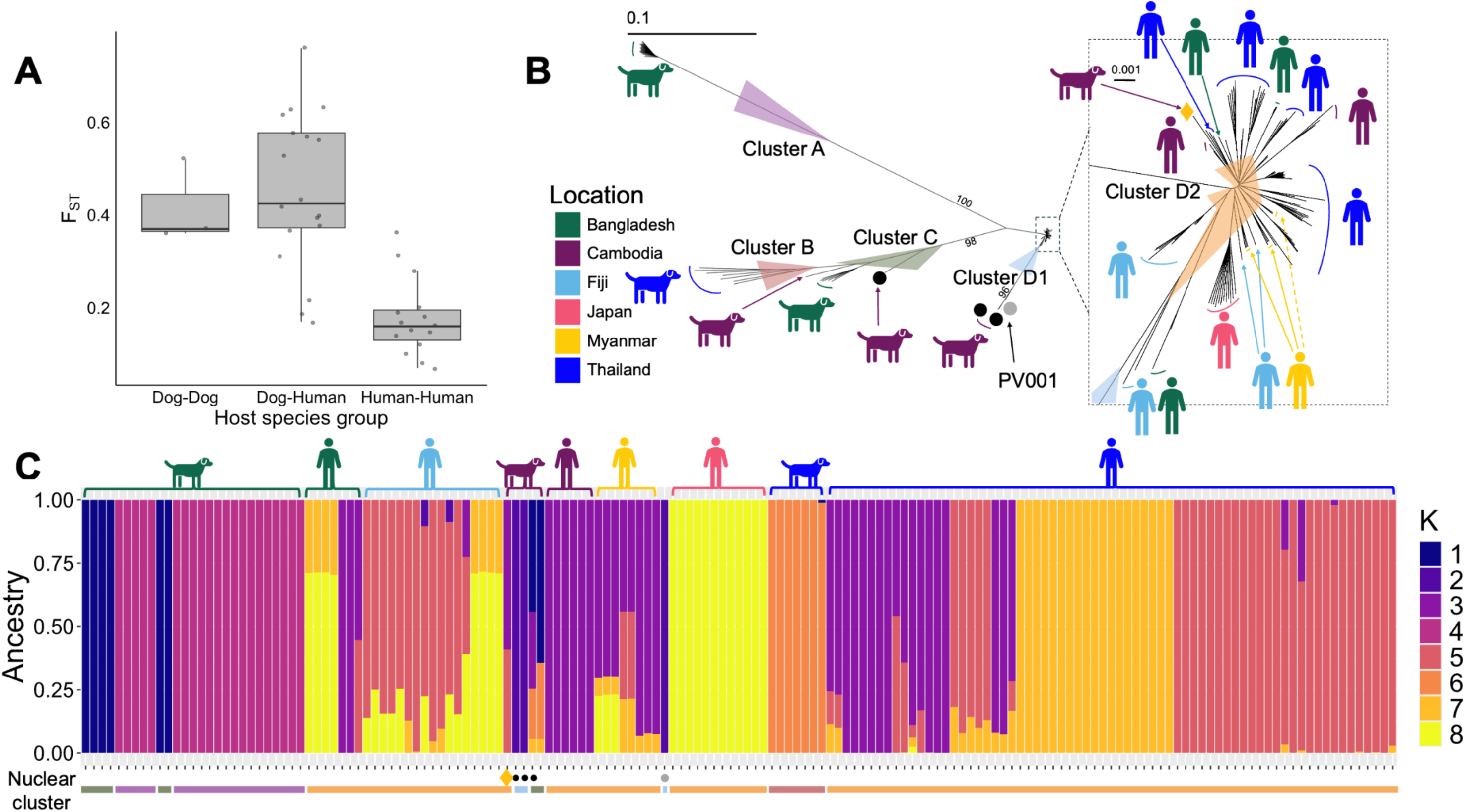
Analysis of the nuclear genome. (A) The F_ST_ values for comparisons between dog-derived, human-derived and human- and dog-derived iL3s; in the box plot the whiskers are 1.5 of the inter-quartile range; (B) A neighbour joining tree of human and dog-derived iL3s, showing clusters A, B, C, D1 and D2; The D2 cluster in the dotted box is expanded to the right; the main tree scale is 1 substitution per 10 bases; this tree with sample IDs is shown in **Figure S8**; bootstrap values >95 % for the main nodes are shown and details of bootstrap values are in **Table S7** (C) Admixture for k = 8, with the clusters from (B) shown below the plot; the iL3 identifiers are shown in **Figure S5.** In A and B, a putative human-to dog cross infection is shown as a yellow diamond; 3 putative introgressed genotypes (KHD19_c01, KHD19_c02, KHD32_c; KHD32_c is in clade C) as black dots, and the *S. stercoralis* reference genome (PV001) as a grey dot. The colour code for country of origin for the dog and people icons is the same in B and C.

We constructed a neighbour joining (NJ) tree of the human- and dog-derived iL3s. This revealed four clades of dog-derived *Strongyloides*, and a single clade of human-derived *Strongyloides*, but which included a single dog-derived iL3 (**Figure 2B**). The clade containing all the human-derived *Strongyloides* generally has short branch lengths, and there is limited separation by the iL3s’ country of origin. This contrasts with the four clades containing only dog-derived iL3s, which are separated on generally long branch lengths. These same 5 groups of human and dog-derived iL3s are evident in a Principal Component Analysis (PCA) of the same data (**Figure S3**). These NJ and PCA results are consistent with the F_ST_ analysis, showing genetic differentiation between human- and dog-derived *Strongyloides*, that the human-derived *Strongyloides* from six countries in Asia are genetically closely related, while the dog-derived iL3s from three countries in Asia are genetically comparatively diverse.

Across the 158 iL3s almost all iL3s from the same host cluster together. The exception is a Bangladesh dog (BDD31) that produced iL3s belonging to two different dog-derived clades, and a person from Fiji (F1) who produced iL3s that occurred in different sub-branches of the clade containing all the human-derived iL3s (**Figure S4**). Conversely, there are two examples where very closely genetically related iL3s came from different people, in both Cambodia (KHH83 and KHH01) and Thailand (THH61 and THH63) (**Figure S4**). These results show that *Strongyloides* genotypes can be widely distributed across Asia, also suggesting geographical movement of them, presumably as their human and dog hosts move.

We looked for evidence of admixture among the iL3s, with ADMIXTURE best supporting k = 8 groups, three of which are specific for human-derived iL3s (though including a single dog-derived iL3s, as the NJ tree, above), and four for dog-derived iL3s (**Figure 2C**). Admixture was more common among human-derived iL3s (52 of 119 = 44%) than dog-derived iL3s (4 of 39 = 10%). ADMIXTURE for k = 5 supports the 5 clades seen in the NJ tree (**Figure S5**).

To check that our sample combination strategy did not affect our results we repeated our analyses removing the 15 combined samples. This showed that the overall NJ tree structure and the F_ST_ values were not affected, but that ADMIXTURE now identified k = 11 groups, and admixture was more common in the dog-derived iL3s (**Figure S6; Table S5**). We sampled multiple iL3s from some hosts, but these may be identical siblings due to the parthenogenetic reproduction of the parasitic females. To check that this scenario did not affect our results we (i) down-sampled to only a single iL3 per host and (ii) used KING to identify identical iL3s from single hosts, which detected 35 that we then removed, and then repeated the F_ST_ and Admixture analyses. This showed no qualitative change to our results, with high F_ST_ values between human- and dog-derived iL3s and among dog-derived iL3s, but low F_ST_ among human-derived iL3s and Admixture groups that predominantly contained only human-derived or dog-derived iL3s (**Table S6**). We also analyzed the iL3s in (i) and (ii), above, using dXY, which measures the absolute genetic divergence between populations (**Table S6**). This also showed no qualitative change to our results, with high dXY values between human- and dog-derived iL3s and among dog-derived iL3s, but low dXY among human-derived iL3s (**Table S6**).

Together, these nuclear genome analyses suggest that human- and dog-derived *Strongyloides* are almost exclusively different parasite populations.

### The mitochondrial genome reveals evidence of cross infection and introgression between human and dog parasites

We constructed a Maximum Likelihood (ML) tree of the assembled mitochondrial genomes of the 158 iL3s. This revealed eight mitochondrial clades, three of which contain only dog-derived iL3s and two only human-derived iL3s (**Figure 3A**). Three clades contain both human and dog-derived iL3s: (i) 63 human-derived iL3s from Bangladesh, Cambodia, Thailand and Myanmar together with a single dog-derived iL3 from Cambodia; (ii) 13 human-derived iL3s from Thailand and Fiji with the dog-derived *S. stercoralis* reference isolate PV001; (iii) 17 human-derived iL3s from Bangladesh, Cambodia, Thailand and Myanmar and 3 dog-derived iL3s from Cambodia.

**Figure 3.**
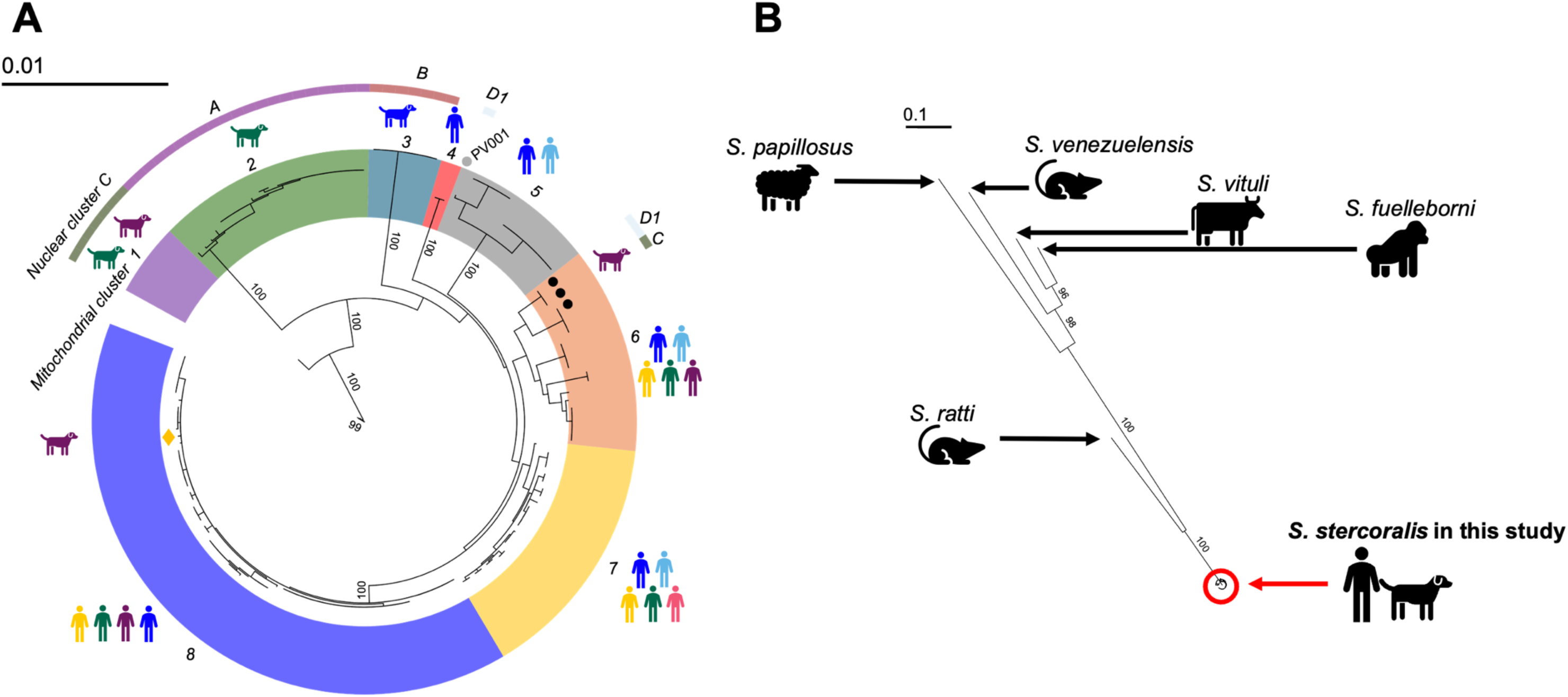
Analysis of the mitochondrial genome. (A) A maximum likelihood tree of human and dog-derived iL3s, showing 8 mitochondrial clusters; the colour code for country of origin for the dog and people icons and the nuclear clusters (A, B, C, D1, D2) are shown as in Figure 2; this tree with sample IDs is shown in **Figure S8**; bootstrap values >95 % for the main nodes are shown and details of bootstrap values are in **Table S7;** (b) A maximum likelihood tree of mitochondrial genomes from this study and of *S. fuelleborni, S. papillosus, S. ratti, S. venezuelensis* and *S. vituli.* The scales in A and B are 1 substitution per 100 and per 10 bases, respectively.

Focussing on the dog-derived iL3s in these majority human-derived mitochondrial clades, and considering the results of the analysis of the nuclear genome (**Figure 2**), there is (a) evidence consistent with contemporary cross-infection of a dog by a human-derived parasite and (b) historical introgression between human and dog-derived iL3s. Specifically, for (a) the single dog-derived iL3 (KHD001_c) among 63 human-derived iL3s belongs to nuclear and mitochondrial clades where all other iL3s are from people. We suggest that this is evidence of a human-derived *S. stercoralis* naturally infecting a dog, thus a reverse zoonotic event. For (b) three dog-derived iL3s (KHD19_c01, KHD19_c02, KHD32_c) belong to a nuclear clade of dog-derived iL3s, but mitochondrial clades of human-derived iL3s, which we suggest that this is evidence of historical introgression between human- and dog-derived *S. stercoralis*. In this scenario we envisage that human- and dog-derived *Strongyloides* co-infected a host, that free-living adult males and females in faeces then mated, and the progeny iL3s then infected a dog, with descendant parasites mating further with dog-derived parasites (**Figure 4**). We also note that KHD32_c is one of the four dog-derived iL3s with evidence of admixture among two human-derived and two dog-derived ADMIXTURE-defined groups (**Figure 2C**), also consist with historical introgression. The *S. stercoralis* PV001 reference genome (isolated from a dog in the USA in the 1980s (24)) similarly appears to be an historically introgressed genotype, belonging to a nuclear clade of Asian dog-derived iL3s, but mitochondrial clades of human derived iL3s.

**Figure 4.**
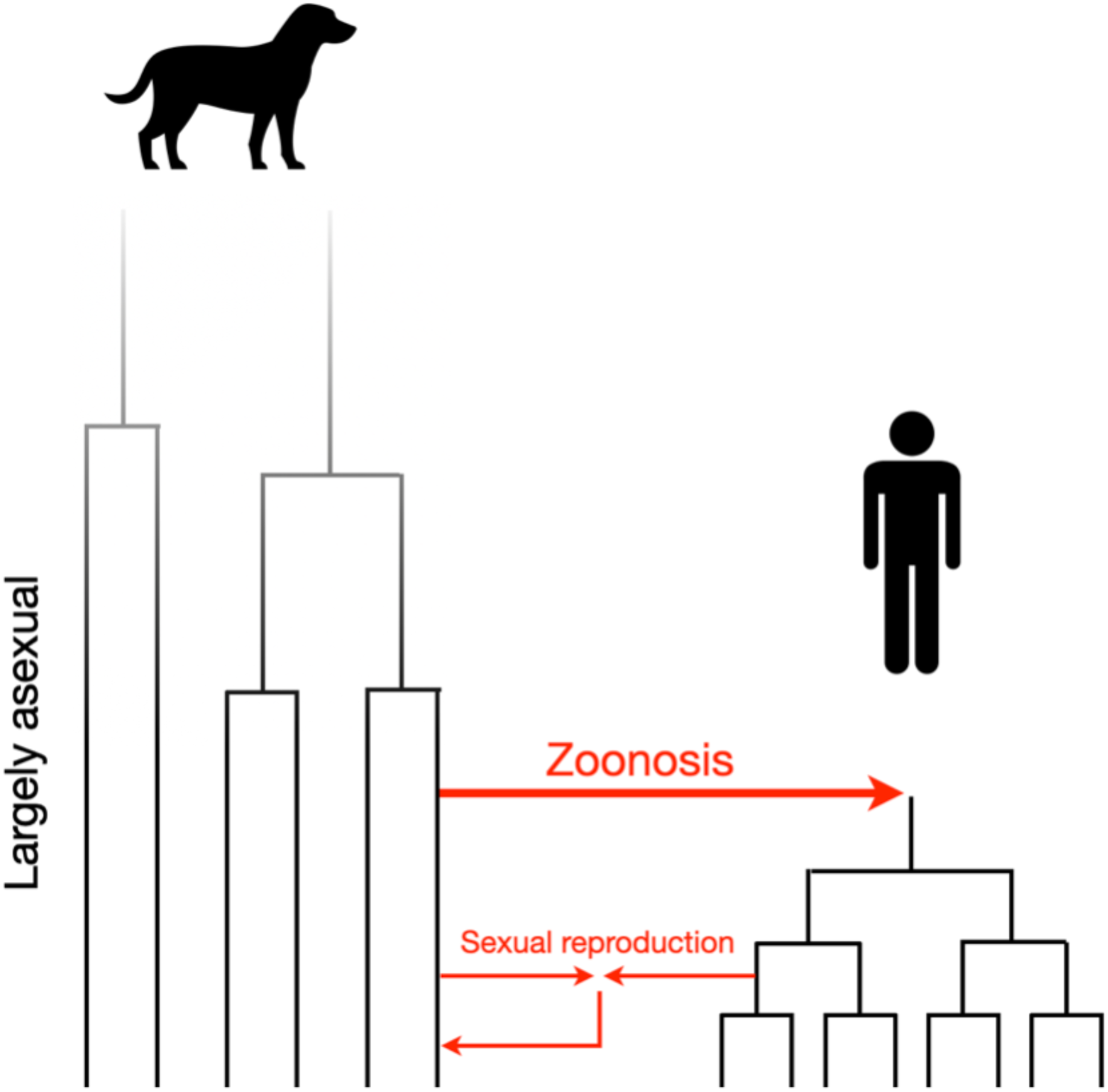
Origin and basis of human- and dog-derived *S. stercoralis.* The hypothesised evolution of *S. stercoralis* from dogs, that zoonotically infected people when dogs were domesticated, with subsequent largely host-specific transmission, but with some introgression between dog- and human-derived parasites.

We calculated the genetic divergence between the mitochondrial genome of human- and dog-derived parasites. Assuming the *C. elegans* mitochondrial mutation rate and two *Strongyloides* generations per year (20) by direct enumeration of substitutions and by a Bayesian-based method we calculated that the last common ancestors of *S. stercoralis* in clades that are shared between people and dogs occurred 5,400 – 8,000 and 9,500 – 12,000 years ago (Bayesian and direct enumeration, for clades 5 and 6, respectively; **Figure 3A**), which is approximately when humans domesticated dogs *c.*10,000-15,000 years ago (25). The last common ancestors of human-derived *S. stercoralis* was 2,000 – 3,400 and 2,000 – 4,000 years ago (Bayesian and direct enumeration for clades 7 and 8, respectively; **Figure 3A**). We therefore propose a model of human *S. stercoralis* infection, originally being a canid parasite from which a lineage infected humans, but that there has been (and continues to be) introgression among these otherwise separate parasite populations (**Figure 4**).

We also sought to understand regions of the genome that might underly the human *vs*. dog infection preference of *S. stercoralis.* We did this by analysing F_ST_ across the genome, with the logic that genomic regions that might generate different host preferences would also be genetically diverged, and so have comparatively high F_ST_ values, differentiated between human- and dog-derived parasites. This showed that there are regions all across the genome with high F_ST_, though particularly on the autosomes rather than the X chromosome (**Figure S7**).

We also wanted to further understand the relationship of these human and dog-derived iL3s to other *Strongyloides* species, to consider whether the human- and dog-derived parasites are themselves different species. To do this we constructed a ML mitochondrial tree of our samples with *S. ratti* and *S. venezuelensis* (both from rats), *S. papillosus* and *S. vituli* (both from ruminants), and *S. fuelleborni* (from primates) (**Figure 3B**). This showed that our *S. stercoralis* iL3s from people and dogs are all very closely related to each other, but quite distinct from other species. We therefore conclude that the iL3s we have sampled from people and dogs are not separate species.

## Discussion

Our whole genome sequencing of *S. stercoralis* from people and dogs across Asia has shown that the human- and dog-derived iL3s are very largely genetically non-overlapping populations. The parasitological interpretation of this is that *Strongyloides* infecting dogs is mainly transmitted among dogs, and *Strongyloides* infecting people is mainly transmitted among people and therefore that dogs are not a major source of zoonotic infection. However, our data also show some exceptions to this. Specifically, we discovered a single dog-derived iL3 (<1% of all 158 iL3s) that is genetically very similar to human-derived *S. stercoralis*. Our parasitological interpretation of this is that a dog was infected by a human-derived iL3s, a reverse zoonosis. There has also been a recent report of a single human-derived worm (of 24 *S. stercoralis* larvae from 8 people) in Bangladesh that genetically clusters with dog-derived parasites, which parasitologically can be interpreted as a zoonotic infection (4).

Further, we and others have identified iL3s where the nuclear and mitochondrial genomes belong to discordant human and dog genetic clades. Specifically, (i) three iL3s (*c*.2% of all 158 iL3s) from dogs with nuclear genomes clustering with those of other dog-derived iL3s, but whose mitochondrial genomes cluster with other human-derived iL3s, (ii) two dog-derived larvae from a single dog with nuclear genomes that clustered with other human-derived larvae, but mitochondrial genomes that clustered with human- and dog-derived samples (4). Our interpretation of this is that there has been introgression between human- and dog-derived *S. stercoralis*. We also found evidence of contemporary cross-infection between parasites from humans and dogs, which might indicate that the opportunity for introgression continues. An alternative hypothesis for these observations is incomplete lineage sorting. That is, that nuclear and mitochondrial genetic diversity present in ancestral *S. stercoralis* has largely, but incompletely, separated into human- and dog-derived parasites. Introgression has been inferred in *Ascaris* infecting people and pigs (12, 13), and our results therefore are potentially a second example of human and animal nematode parasite introgression.

The dog-derived *S. stercoralis* has long nuclear tree branch lengths and rarer evidence of admixture suggesting that it may have a preponderance of direct, asexual reproduction in its history. Further, its population genetic structure may consist of an assemblage of lineages, as has been observed for *S. ratti* in the UK (20). It is conceivable that such different lineages have evolved different host preferences including some for infection of people.

We estimated the divergence time of human- and dog-derived *S. stercoralis* as 5,400 – 12,000 years ago, which is approximately the same time as human domestication of dogs. We therefore suggest that (i) ancestral *S. stercoralis* was a parasite of wolves that evolved into a parasite of dogs, which then infected people, and (ii) since dog domestication, *S. stercoralis* in dogs and people have become genetically divergent, but not reproductively separated. Our results show that dog-derived *S. stercoralis* are much more genetically diverse than human-derived *S. stercoralis*, which could be due to only limited historical transmission of dog parasites to people, which has effectively bottlenecked *S. stercoralis* infecting people.

Our results raise the question of whether the largely separate human and dog transmission cycles that we infer are because of different and diverging host preference of human and dog *S. stercoralis* and / or because of environmental settings that favour person-to-person and separate dog-to-dog transmission. Anecdotal observation at our study sites is that there is human, dog (and other animal) faecal contamination, and close proximity between people, dogs and other animals. Such settings would suggest that there are, in principle, ample opportunities for cross-host species transmission, as much as within-host species transmission. Based on this we therefore suggest that the largely different transmission cycles that we infer are because of different host preference of the parasites. Specifically, that human-derived *S. stercoralis* have a comparatively higher fitness when infecting people than dogs and, *vice versa*, that dog-derived *S. stercoralis* have a comparatively higher fitness when infecting dogs.

Many, perhaps most, parasitic nematodes appear to have a high host specificity, that is that they can successfully infect and reproduce in a single (or small number of) host species, but infect less well (or not at all) other host species. Despite the importance of host specificity of parasitic nematodes there has been little investigation of this, usually requiring host cross-infection experiments. For *Strongyloides*, apart from the human-dog cross infection experiments (above), *S. ratti* and *S. venezuelensis* (both parasites of rats) have been used to infect mice, where high infective doses are required to establish an infection, and then the infection is abbreviated, compared with infections in laboratory rats (26). Genomic analyses has the potential to be used to investigate host specificity, for example, by comparing parasites with different host specificities and preferences to identify genomic regions that differ between them, with such regions being putatively involved in determining host specificity. Our sliding-window F_ST_ analysis across the genome showed that there were many regions that differ between human- and dog-derived parasites, suggesting a complex genetic basis of host specificity. It would be interesting to investigate this further, but a key first step would be to generate high quality genome assembles of human- and dog-derived parasites.

The lab-maintained *S. stercoralis* isolate (PV001) was obtained from a dog in the USA in 1984 (24). Our analysis suggests that it too is an introgressed genotype between human- and dog-derived *S. stercoralis*. Because PV001’s nuclear genome is more similar to dog-derived iL3s than to human-derived iL3s, this must question the validity of PV001 as a laboratory model of human-infecting *S. stercoralis.* We therefore suggest that it would be desirable to newly isolate *S. stercoralis* from people and maintain it in dogs, if indeed this is possible.

Despite the largely separate human and dog transmission cycles of *S. stercoralis* that we infer, these parasites are very closely related, compared with other *Strongyloides* species. The name *S. stercoralis* is used for parasites from people and dogs, but the name *S. canis* was also created for parasites from dogs (27). This species name was created based on difficulty in human to dog infection, and different geographical ranges, but the genetic results presented here do not support its differentiation from *S. stercoralis*. More generally, *Strongyloides* species descriptions are principally morphological though host species has also been part of some descriptions. These species descriptions are challenging because of the limited morphological taxonomic characters available, and that some of the more useful ones characters are present on the within-host parasitic stages, and so are more rarely available for inspection (and very rarely so for those infecting people). Further, the use of host species to identify *Strongyloide*s species is inherently circular when trying to understand the host range of different *Strongyloides* species.

Given the close genetic similarity between *S. stercoralis* from people and dogs, there is the question of whether these should be considered separate species. We suggest that such a designation is premature, for three reasons. One, while there are genetic differences among the human-derived and dog-derived parasites, they are very similar compared with the differences with other *Strongyloides* species (**Figure 3B**). Two, there is evidence of hybridization between human- and dog-derived parasites, though, further work is needed to measure the frequency of this. Three, our study only pertains to Asia, but *S. stercoralis* occurs throughout the tropics and the genetic relationship between human- and dog-derived parasites in Africa, Latin America and Australasia remains to be investigated. We therefore suggest, pending further study, that the name *S. stercoralis* continues to be used for both human and dog-derived parasites, though the host species source of any parasites is always presented.

Our study has limitations. One, while we studied a total of 158 iL3s, a larger sample size would give a better estimate of cross-infection and introgression events, which we only detected rarely. Two, we sampled multiple iL3s from some hosts, which may include identical siblings, though we accounted for this by down-sampling, identifying and then excluding identical siblings, and using alternate measures of genetic differentiation, and found that our results and conclusions were robust to these alternative analyses. Three, our work has focussed on *S. stercoralis* in Asia, and so it remains unclear whether there is a similar biology and genetics of *S. stercoralis* in other endemic regions, particularly Africa and South America. Four, our work has used the *S. stercoralis* reference genome, but given the genetic divergence between human- and dog-derived *S. stercoralis* then better genetic resolution would be obtained by using alternative reference genomes.

## Materials and Methods

### Parasite sampling

Faeces were sampled from people and dogs in: (i) Chittagong, Bangladesh in January 2020 and January 2023; (ii) Preah Vihear, Cambodia in August 2023 and (iii) Khon Kaen Thailand in July 2022 (**Table S1**), and cultured to obtain iL3s (**Supplementary Methods**).

We also obtained samples from three Fijian born migrants to the UK. Following testing at the Liverpool School of Tropical Medicine they were found to be infected with *S. stercoralis*. These individuals have no UK-based risk of *Strongyloides* infection and therefore their infections are thought to likely represent *S. stercoralis* acquired in Fiji (28).

The Bangladesh, Cambodia and Thailand-derived samples iL3s are identified as: country abbreviation + host species + host ID + order of collection. Country abbreviations: BD, Bangladesh; KH, Cambodia; TH, Thailand. Host abbreviations: H, human; D, Dog. Thus iL3s “THD141_01” is the first iL3 collected from a dog 141 in Thailand (**Tables S1 and S3**). Fijian-derived samples are numbered using the F country code, but omit the host code.

### DNA sequencing and quality control

DNA was prepared from individual iL3s as described by (20), sequenced, and aligned to the *S. stercoralis* reference genome (**Supplementary Methods**). After genome alignment, we quality controlled (QC) the data using two criteria for inclusion in analyses: (i) ≥20 average depth of coverage across the whole genome and (ii) ≥80% genome coverage.

### Combining sample sequences

For some samples from Cambodia, Fiji and Thailand the sequence reads from individual iL3s did not meet the QC inclusion threshold, or only a single iL3s sample did. Here we used a sample combination strategy by identifying SNPs (as below) and using these to construct a neighbour joining (NJ) tree (as below) and identified individual iL3s that (i) originated from the same host and (ii) clustered within identical nodes on the tree, which we then combined into single dataset to pass the QC criteria, and then included them in our analyses (**Figure S2; Table S2**). We also constructed a NJ tree of mitochondrial SNPs of all samples, and confirmed that the samples we combined were within the same nodes. These sequence samples use the naming convention described above, but with “c”, instead of order of larval collection.

### Additional sequence data

To our own data we added previously published *S. stercoralis* whole genome data: (i) 3 iL3s from people in Myanmar and (ii) 6 iL3s from people in Japan, both at NCBI PRJDB5112 (**Table S3**), with QC as above. These samples have the codes MyHTB for Myanmar and RK for Japan and then are followed by individual sample numbers.

### Bioinformatics – nuclear genome

For Bangladesh, Cambodia, Thailand, Fiji, Myanmar, and Japan iL3 from QC-passed sequence data we detected SNPs compared to the *S. stercoralis* reference genome using BCFtools with the recommended multiallelic calling option. SNPs were kept if they had: (i) a QUAL score above 20, (ii) a Mapping Quality score above 20, (iii) a Genotype Quality score above 20, (iv) a mean depth above the tenth percentile and below the ninetieth percentile of all SNPs across all samples, (v) a missingness below 1%, and (vi) a minor allele frequency greater than 0.01. SNP sites with nucleotides that were identical among all samples but different from the reference genome were removed.

From these data we calculated a distance matrix for all pairwise combination of iL3s, isolation by distance, F_ST_, constructed a NJ tree, used Principal Component Analyses (PCA), and measured admixture (**Supplementary Methods**). We also determined F_ST_ values between human- and dog-revied parasites across the genome (**Supplementary Methods**).

### Bioinformatics – mitochondrial genome

We mapped sequence reads to the *S. stercoralis* mitochondrial reference genome (NCBI Reference Sequence: NC 028624.1) using Bowtie2 and called and filtered SNPs as above. We assembled mitochondrial genomes and aligned them in two sets: (i) all the iL3 data referred to above and (ii) all other *Strongyloides* mitochondrial genomes with (i) and constructed maximum likelihood (ML) trees, from which we estimated the divergence time between human- and dog-derived *Strongyloides* (**Supplementary Methods**). For the ML trees node support was determined from 500 bootstrap replicates, and is reported as percentages.

### Ethics

This work was approved by the University of Chittagong Ethical Review Board, reference CU SCE210001; the Khon Kaen University Ethics Committee for Human Research, reference HE641605; the Institutional Animal Care and Use Committee of Khon Kaen University, reference 660201.2.11/495 (60); the National Ethics Committee for Health Research, Ministry of Health, Cambodia, reference No. 145NECHR, and the Central University Research Ethics Committee, University of Liverpool, references 10936 and 12514. All human- and dog-derived samples were anonymised, and sequencing reads belonging to the host were removed.

### Data deposition

Genome alignment data (stored in bam files) for the 158 samples used in this study are deposited at the European Nucleotide Archive (https://www.ebi.ac.uk/ena/browser/home), with study accession number of PRJEB80566.

## Acknowledgments

We would like to thank: in Bangladesh, Israt Jahan Shonom, Mahmudul Hasan Limon, Mohini Das, Doshor Rahman, Preity Farah Islam, Emu Esrat Jahan, Sartaz Afrin, Trayee Biswas, Syoda Fariha Tabassum, Umme Humayra, Riddita Chakma; Integrated Development Foundation and Sumon Sarkar and Tamal Sacmo; in Cambodia, Chethna Ten, Chanmakara Mean, Molyden Vann; in Thailand, Petcharat Chompo, Shih Keng Loong, Morsid Andityas, Sangduan Wannachart; Katie O’Brian for sharing her Karken2 customised database; the staff of the Centre for Genomic Research for their help with DNA sequence analysis. YL was supported by the China Scholarship Council, and this work was supported by University of Liverpool ODA Research Seed Fund and by the University of Liverpool.

## Supplementary Methods

### Parasite Sampling

For human faeces, volunteers provided faecal samples in containers given to them. For dog faeces, in Thailand and Cambodia dogs were restrained by their owner, the dogs given a saline enema by a veterinary surgeon, then followed until the dogs defecated, when the faeces were collected. In Bangladesh, dog faeces were collected from the ground, taking care to only collect the upper part of the faecal mass.

Human and dog faecal samples were individually cultured as (1), and maintained at local ambient temperature. After 72 hours *Strongyloides* iL3s were collected, washed once in 1% w/v sodium dodecyl sulphate, twice in distilled water, then transferred individually with 10 μL of distilled water (Bangladesh and Thailand) or absolute ethanol (Cambodia) into a microcentrifuge tube; samples in water were stored at −80 °C; samples in ethanol were stored at ambient temperature.

### DNA sequencing and quality control

DNA lysates were purified using AMPure XP beads, and 300 bp libraries constructed with the NEBNext Ultra II FS Kit using the one tenth, reduced volume protocol. The libraries were sequenced on the Illumina NovaSeq platform generating 2x150 bp reads. Sequencing reads were trimmed of adapter sequences using Cutadapt version 1.2.1 (2), and further trimmed using Sickle version 1.2 (3) with a minimum window quality score of 20; reads ≤15 bp after trimming were removed.

Sequencing reads were processed through Kraken2 (4), using a customised database of the default ‘Standard’ database with (i) four *Strongyloides* species (*S. ratti, S. papillosus, S. stercoralis, S. venezuelensis*) (5) and (ii) other parasitic nematodes (6). *S. stercoralis*-identified reads were then aligned to the *S. stercoralis* reference genome (GenBank ID: GCA 029582065.1) using mapper Bowtie2 version 2.4.5 (7) with default settings, and SAMtools version 1.9 (8) then removed unmapped reads.

### Bioinformatics – nuclear genome

Using the filtered SNPs we calculated a genetic distance matrix for all pairwise combination of iL3s using filtered SNPs in TASSEL version 5.0 (9), and genetic distances were then plotted against geographical distances using the ggplot2 package in R, with the significance of the relationship evaluated with a Mantel test.

To investigate the genetic relationship between iL3s from people and dogs we calculated the pairwise fixation index (F_ST_) values among all samples using VCFtools after (10). We made three pairwise comparisons: (i) Dog-Dog, (ii) Human-Human, (iii) Human-Dog. A box plot of these F_ST_ values among these groups was generated using the ggplot2 package in R. We also calculated the human-dog F_ST_ values in non-overlapping 50 kb windows across the genome. Additionally, to account for potential presence of identical siblings among iL3s sampled from individual hosts, we calculated the average nucleotide divergence (d_XY_) for the same three pairwise comparisons using pixy v2.0.0 (11).

We also constructed a Neighbour Joining (NJ) tree of all the iL3s based on the pairwise genetic distance among samples using TASSEL 5, and generated bootstrap values from 1,000 replicates using R with vcfR (12) and ape (13) packages. The resulting tree file was then visualised with iTOL (14). We used Principal Component Analyses (PCA) of the same data performed with PLINK 1.9 (15), which was processed and visualised in R, using tidyverse and ggplot2 packages.

We performed an admixture analysis using ADMIXTURE (16) for K values from 2 to 13, and determined the optimal K value by that with the lowest cross-validation error in R.

We calculated kinship coefficients between individual iL3s using KING v2.3.2 (17) to identify genetically identical siblings among IL3s sampled from individual hosts. Specifically, PLINK-generated BED files were used as the input. Two output files were produced: kin0, which contains kinship coefficients between iL3s from different hosts; and kin, which contains kinship coefficients between iL3s from the same host. Both were visualised as a heatmap using the ComplexHeatmap (18) package in R.

### Bioinformatics – mitochondrial genome

We assembled mitochondrial genomes of each iL3 and combined samples using the ‘consensus’ function of BCFtools. For heterozygous genotypes the reference allele was selected for the consensus sequence.

We aligned mitochondrial genomes using MAFFT v7.310 (19) in two sets: (i) all the iL3 data referred to above and (ii) all other *Strongyloides* mitochondrial genomes; specifically, *S. fuelleborni* (GenBank OL505577.1), *S. papillosus* (NC 028622.1), *S. ratti* (NC 028623.1), *S. venezuelensis* (NC_028229.1)*, S. vituli* (NC 066507.1) and with (i). We constructed a maximum likelihood (ML) tree using RaxML version 8.2.12 (20), using the General Time Reversible model with 500 bootstraps and visualised in iTOL.

To estimate the divergence time between human- and dog-derived *Strongyloides* from the ML data we (i) counted the number of substitutions per site since the last relevant common ancestor and (ii) used BEAST that uses a Bayesian approach to date phylogenies (21). In these calculations we assumed the *C. elegans* mitochondrial mutation rate of 1.05 x 10^-7^ per site per generation (22), and assumed two *Strongyloides* generations per year, after (23).

---

**Figure S1.**
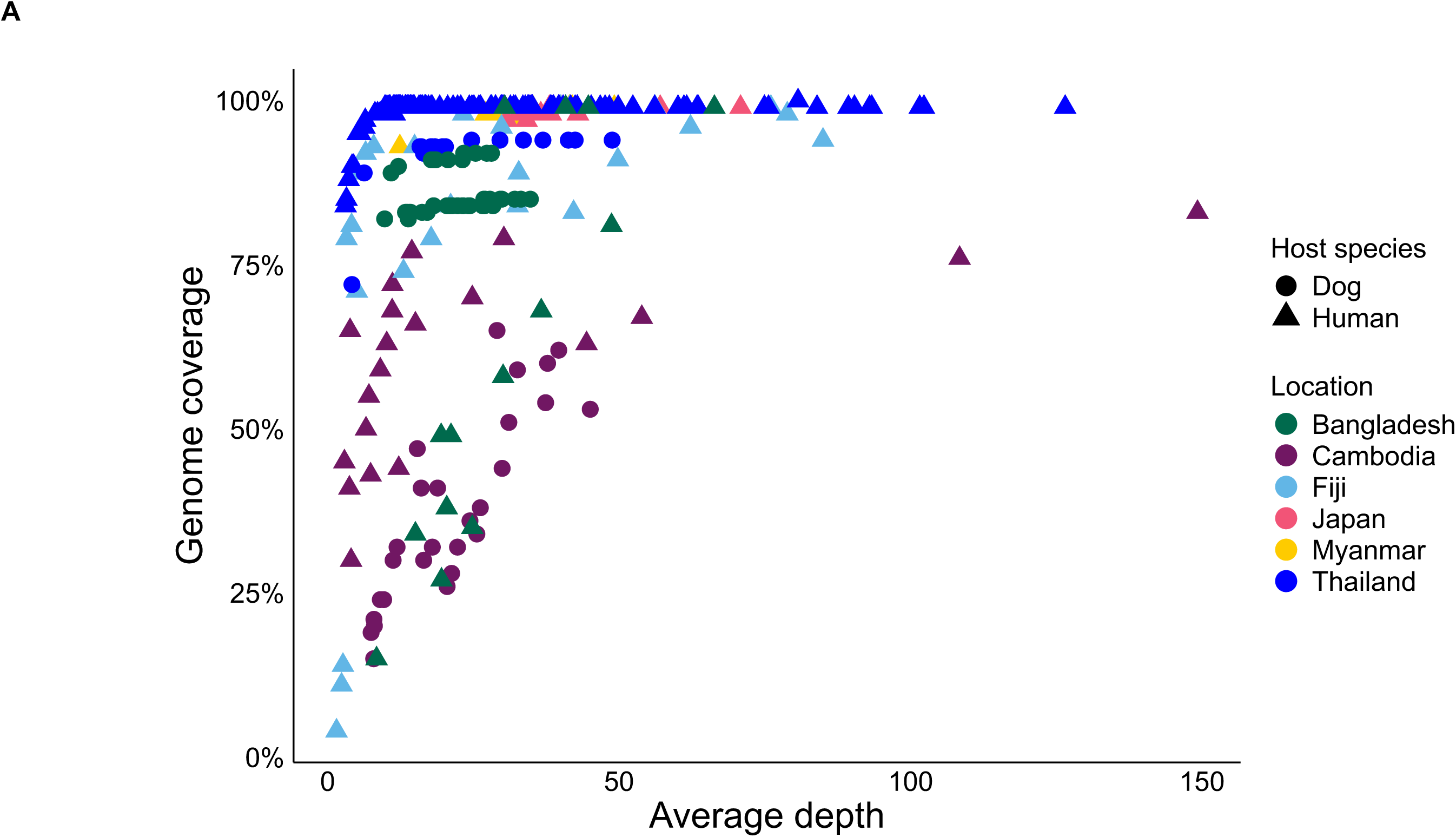
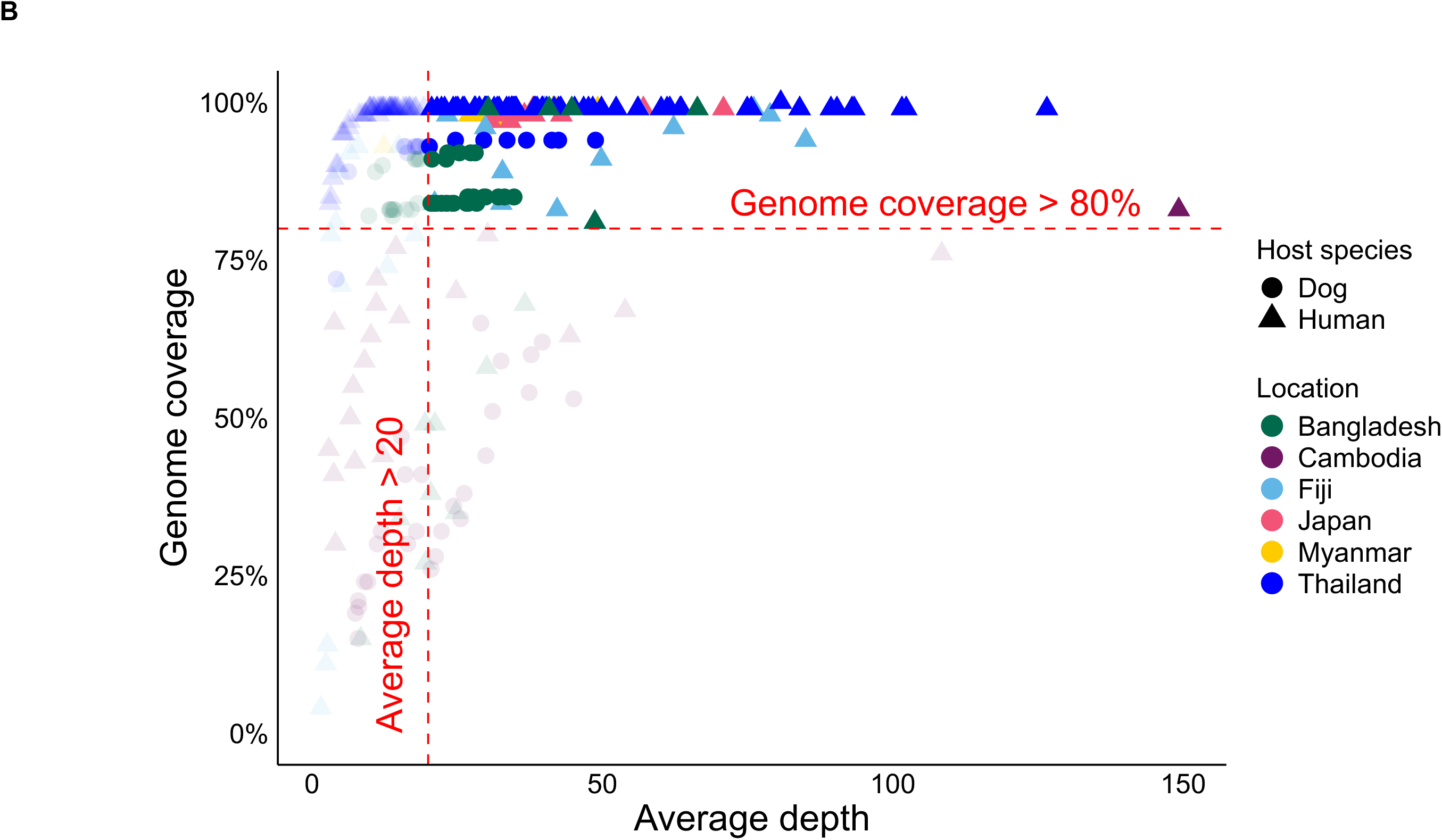
Alignment of sequence reads to the *S. stercoralis* reference genome. The average read depth and genome coverage for (A) all 298 iL3s and (B) for the 143 that passed our quality control criteria. The country and host origin of samples is shown by colour and shape.

**Figure S2.**
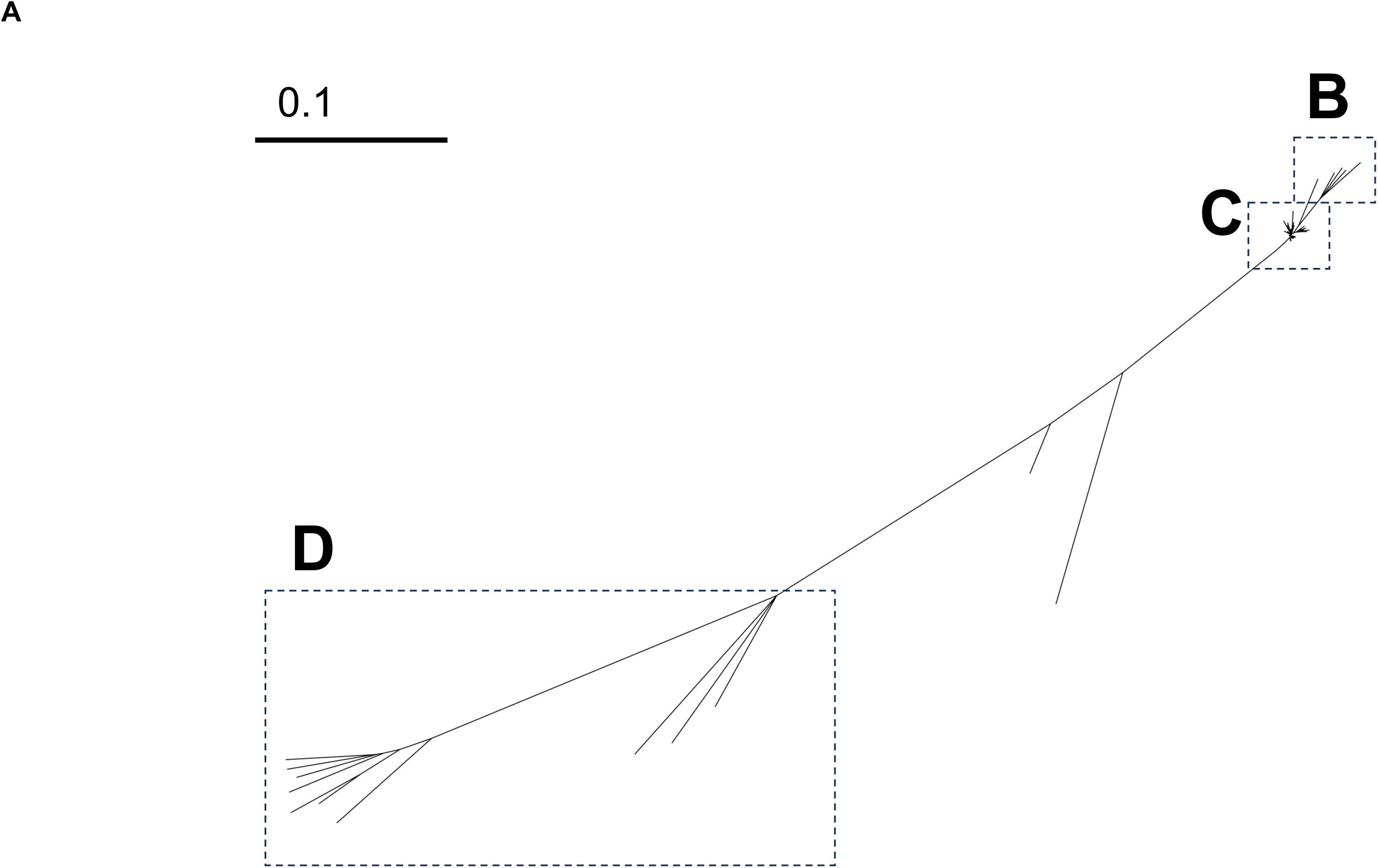
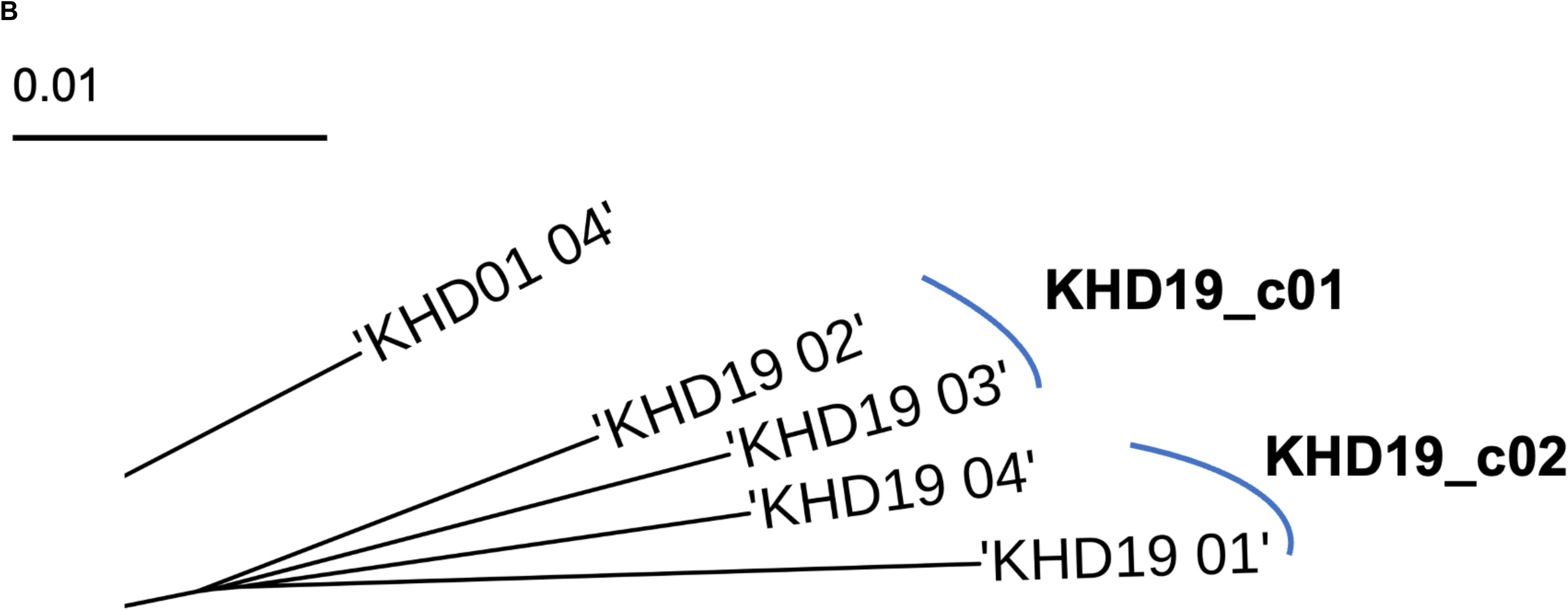
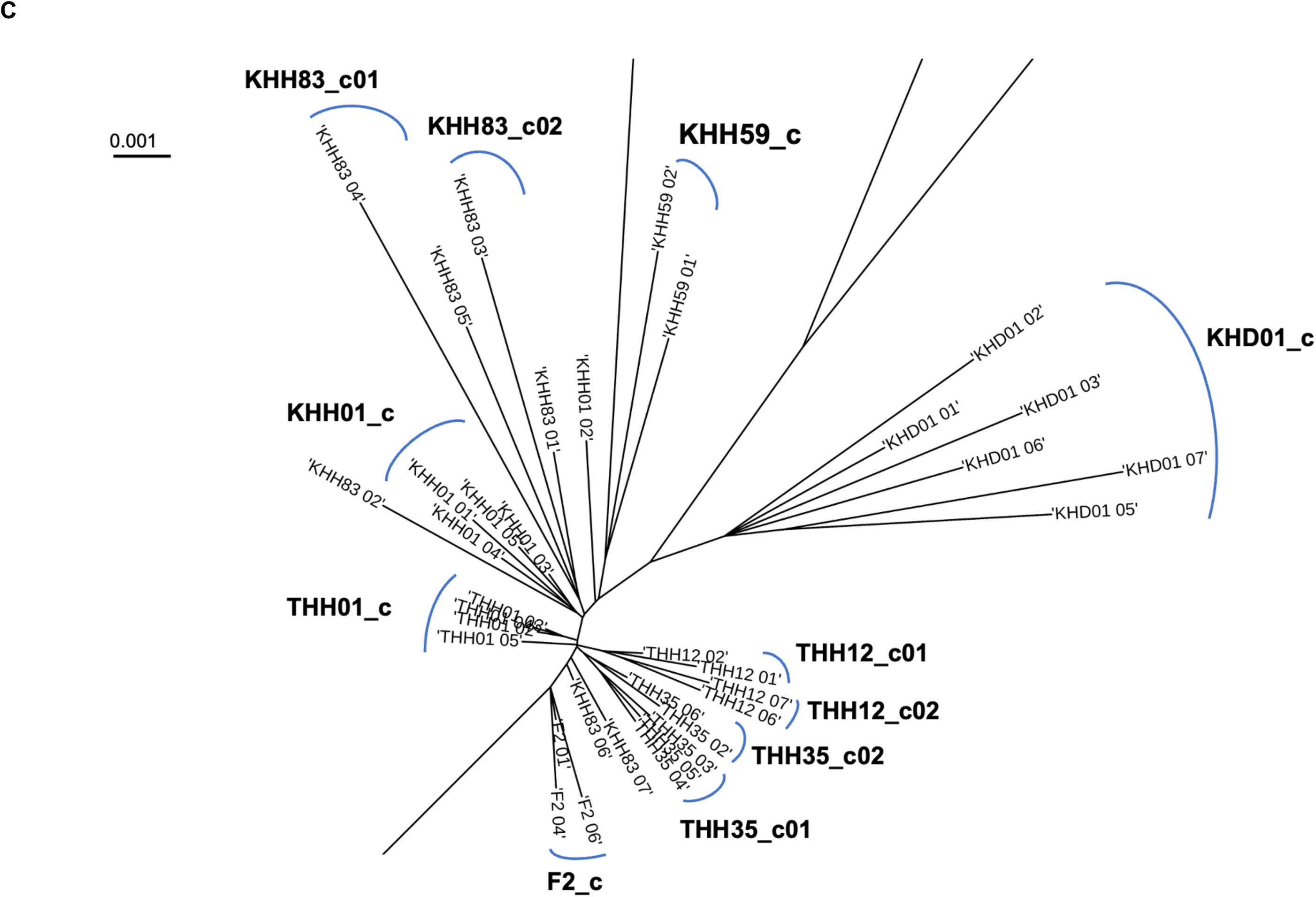
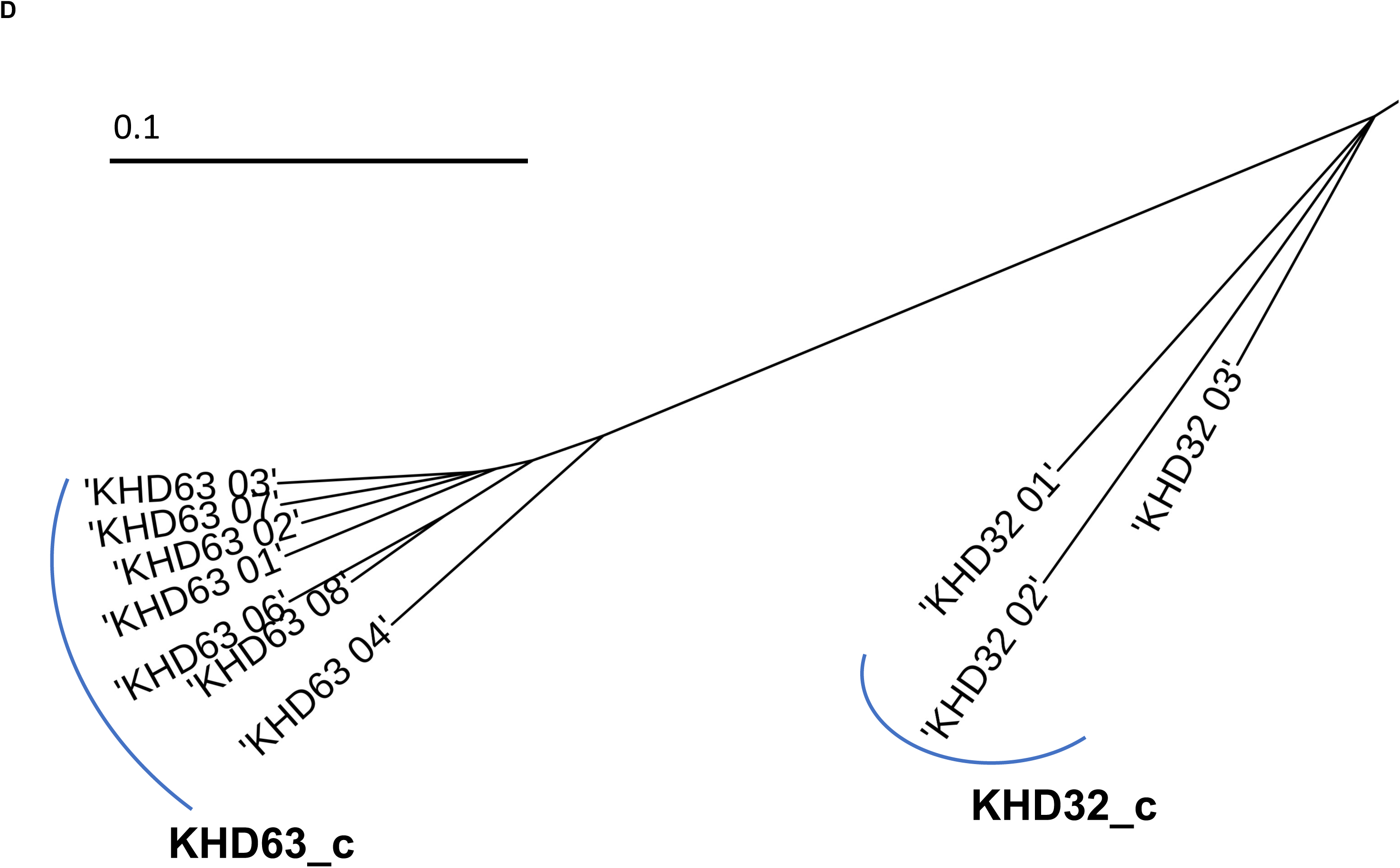
Combined sequence samples. (A) Neighbour joining tree of 56 iL3s where the combination strategy was used. The details in areas **B, C, D** marked by dotted boxes in (A) are shown in (B), (C) and (D), respectively. Samples derived from the same host and within the same nodes are outlined by the blue curves. The scales are 1 substitution per 10 bases in (A) and (D), 1 substitution per 100 bases in (B), and 1 substitution per 1000 bases in (C).

**Figure S3.**
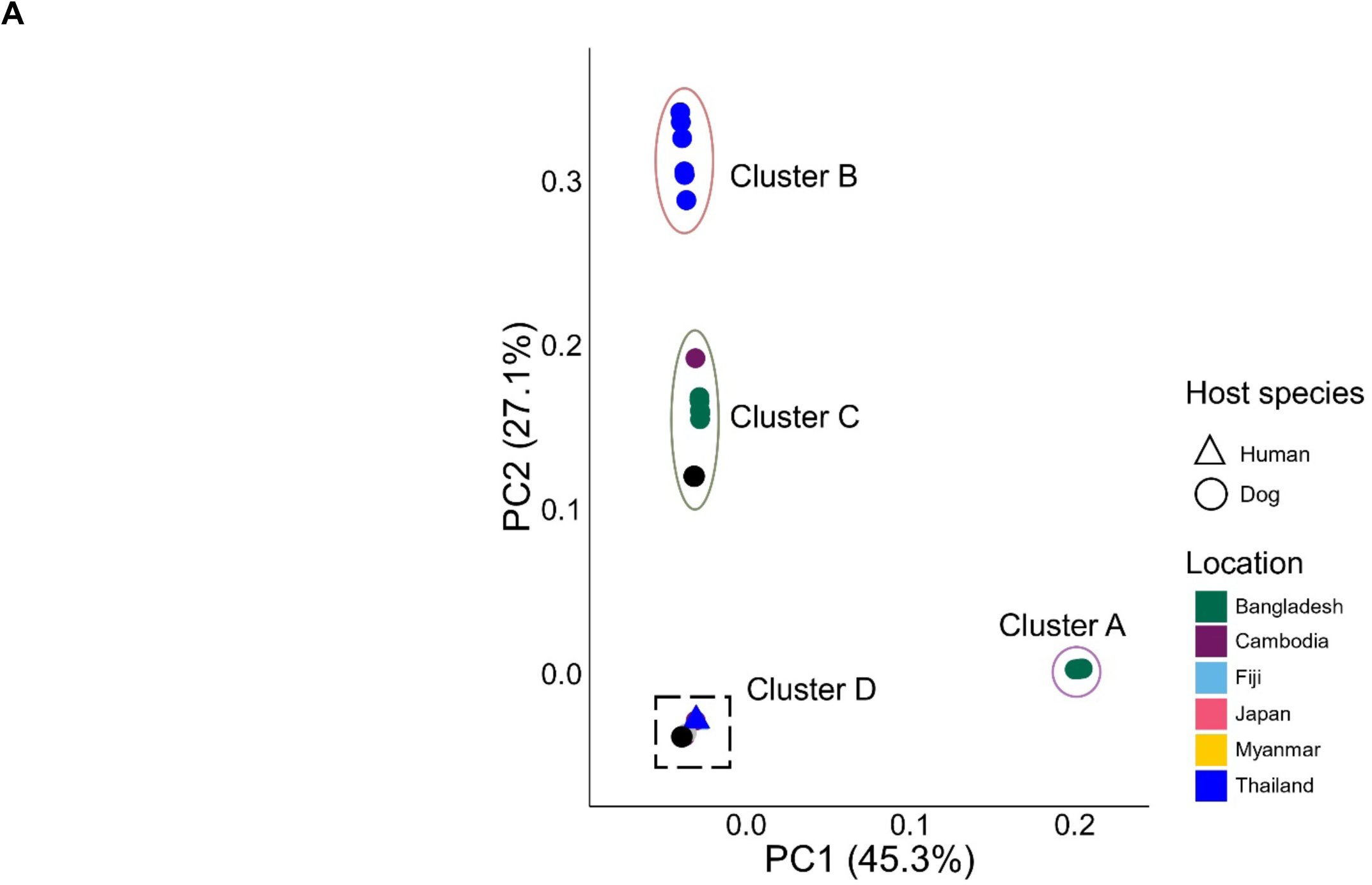
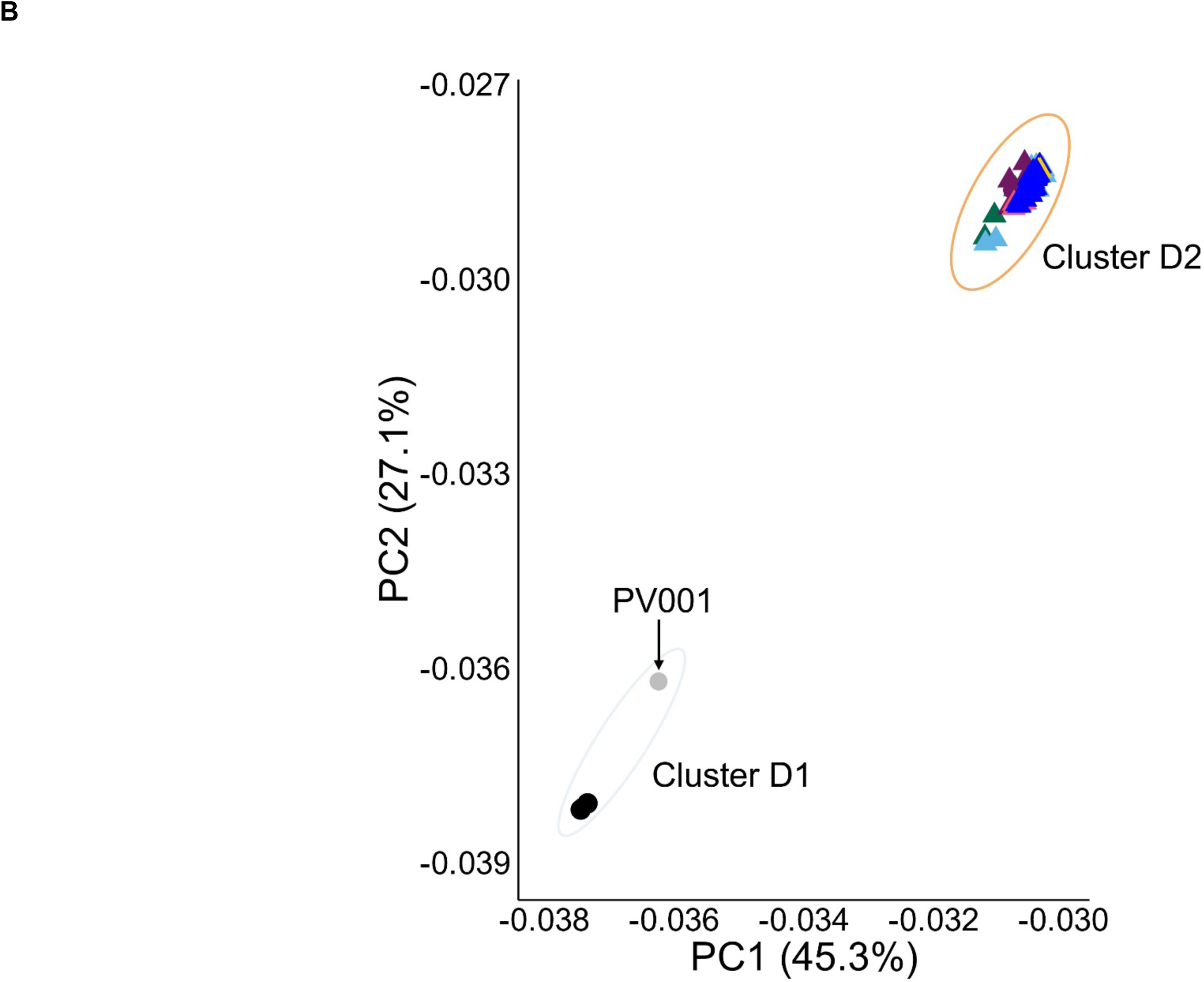
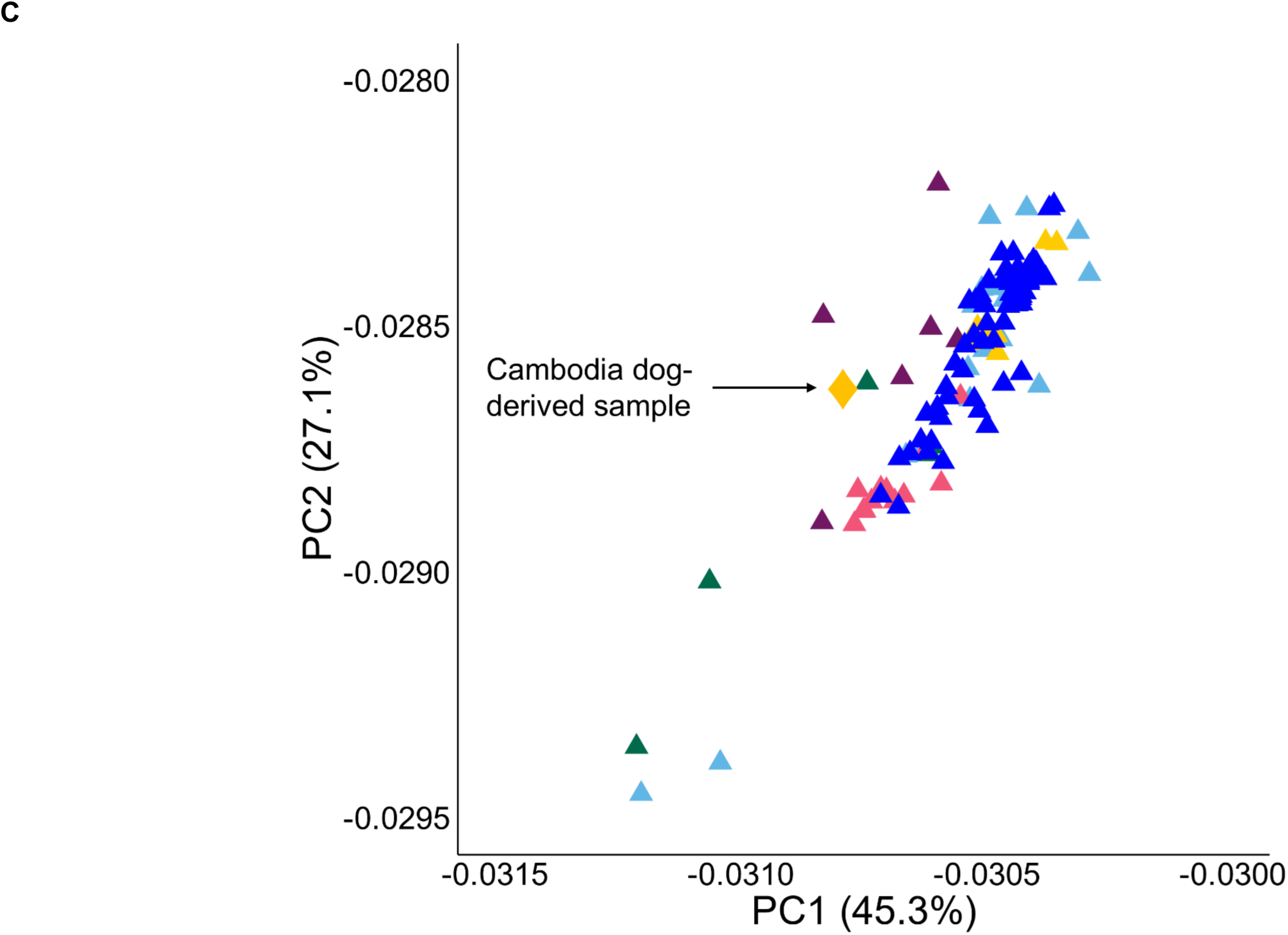
Principal Component Analysis. Plots of Principal Components (PCs) 1 and 2 of (A) all 159 iL3s, (B) those within cluster D, (C) those within cluster D2. The iL3s are shaped and coloured by host species and geographical location. The iL3s are grouped by clusters A, B, C, D, D1 and D2, which are as main text Figure 2B. PCs 1 and 2 account for 45.3 and 27.1 % of the variance, respectively. The grey and black dots in (B) indicates the *S. stercoralis* reference PV001 and 3 putative introgressed genotypes, respectively; the yellow diamond in (C) indicates the putative human-to-dog cross infection.

**Figure S4.**
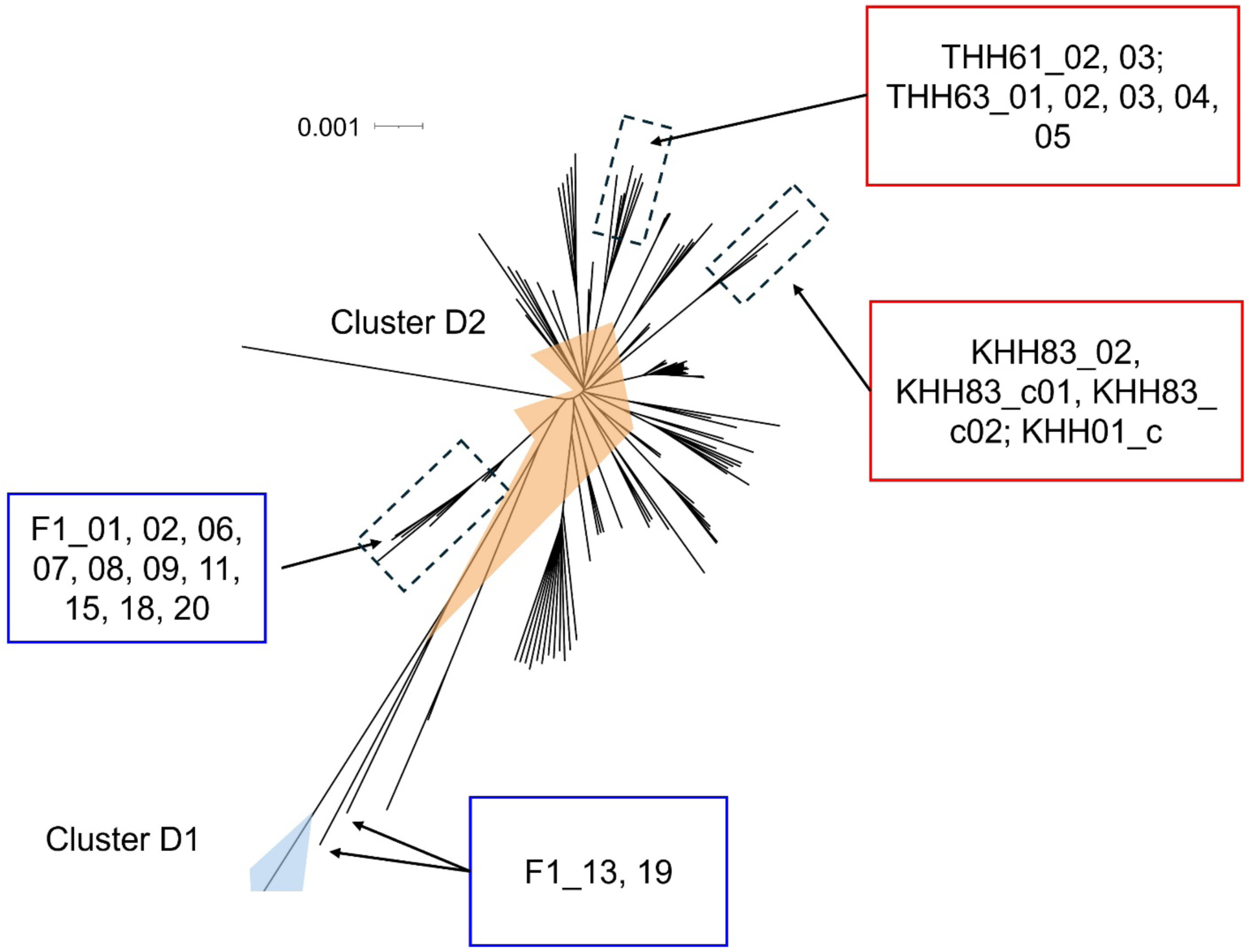
Distribution of genotypes among hosts. Details of cluster D1 and D2 of the neighbour joining tree in main text Figure 2B, showing (i) different genotypes co-infecting a host (blue boxes) and (ii) different hosts in which closely related genotypes occur (red boxes).

**Figure S5.**
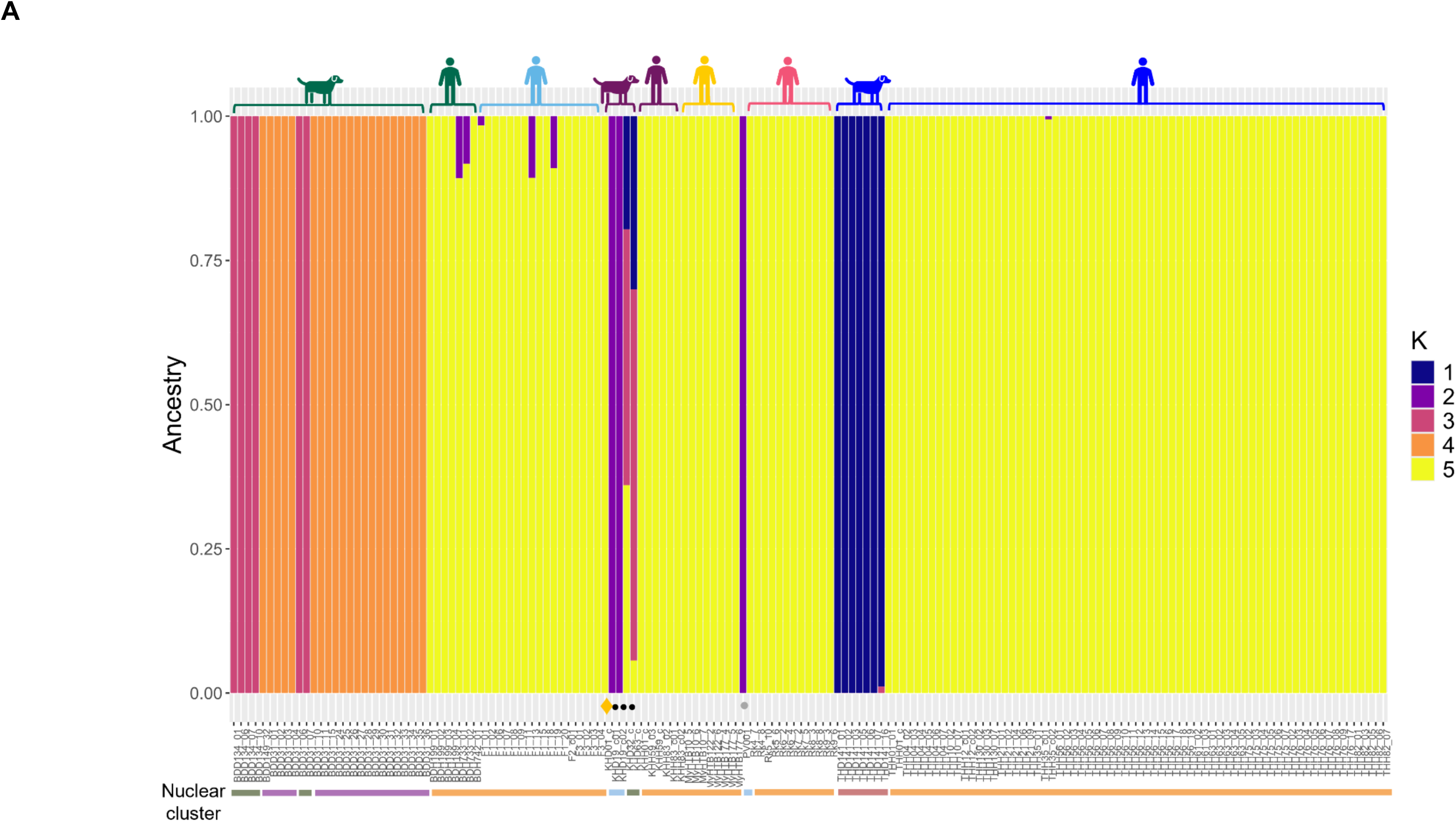
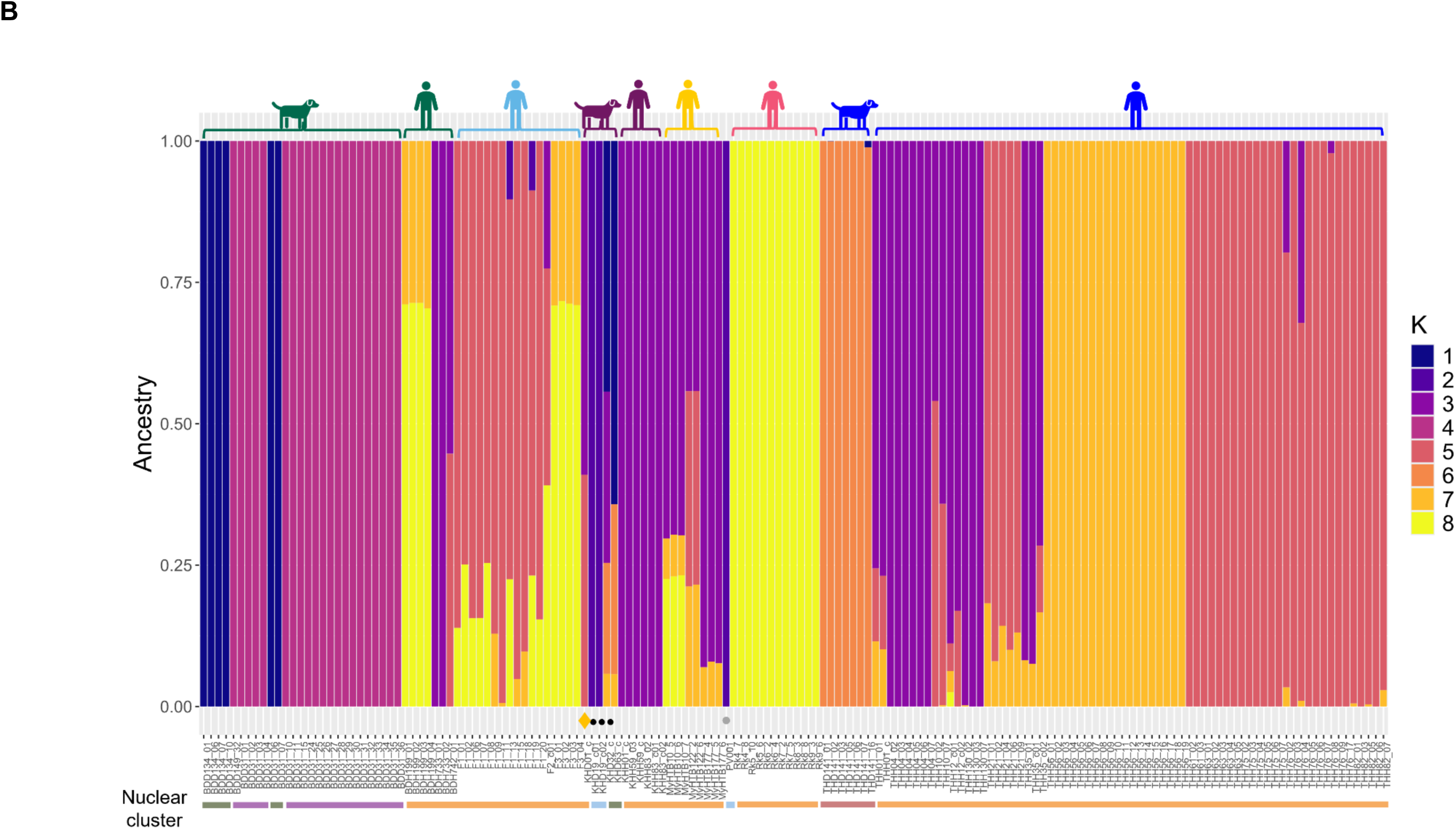
Admixture. (A) Admixture for k = 5. The proportion of admixture and the distribution of genetic ancestries for each iL3, with host species and geographical location of iL3s shown at the top of the plot, and the nuclear clusters of each at the bottom. (B) Admixture for k = 8, as main text Figure 2C, but showing the individual iL3 identifiers.

**Figure S6.**
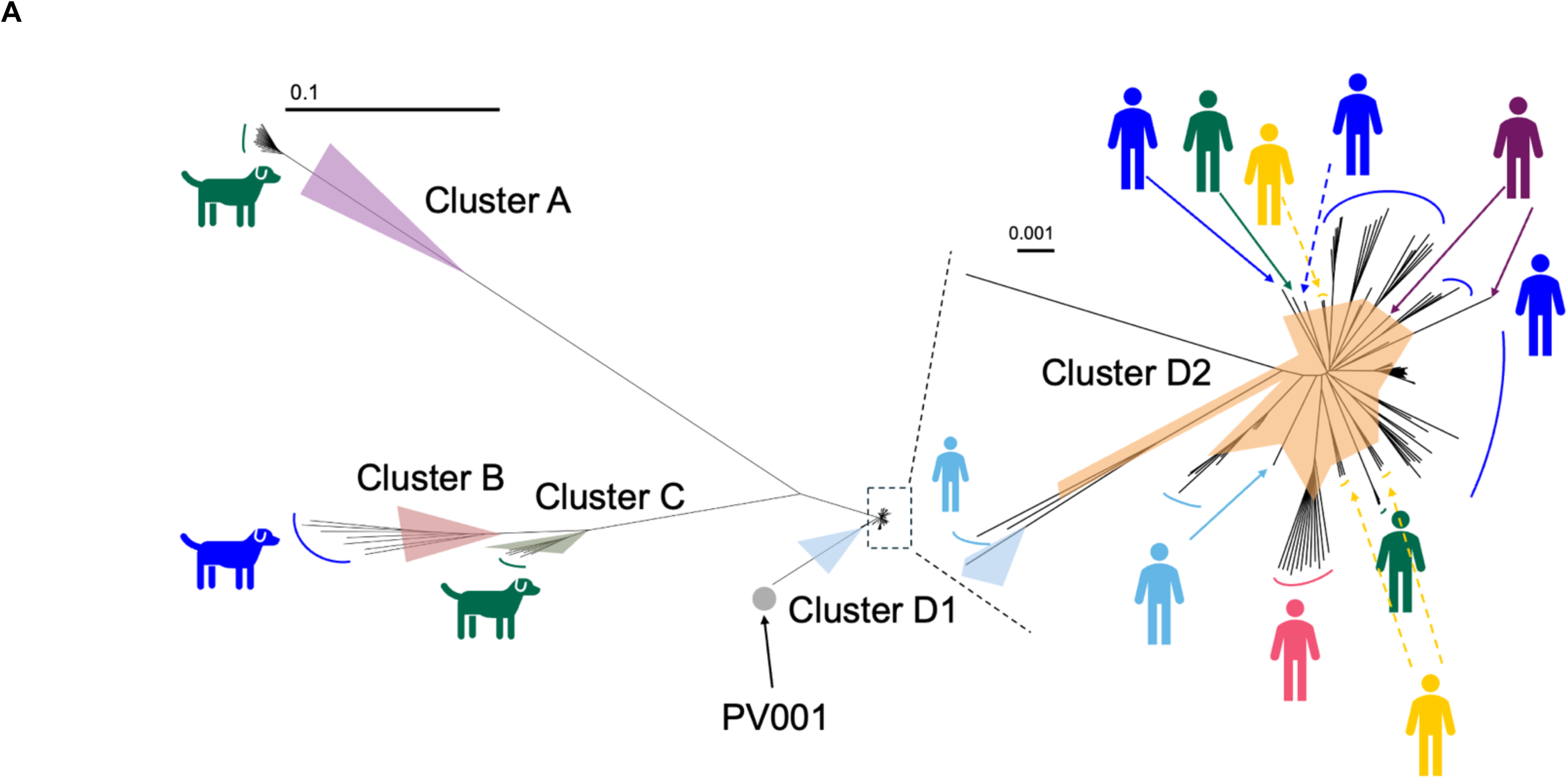
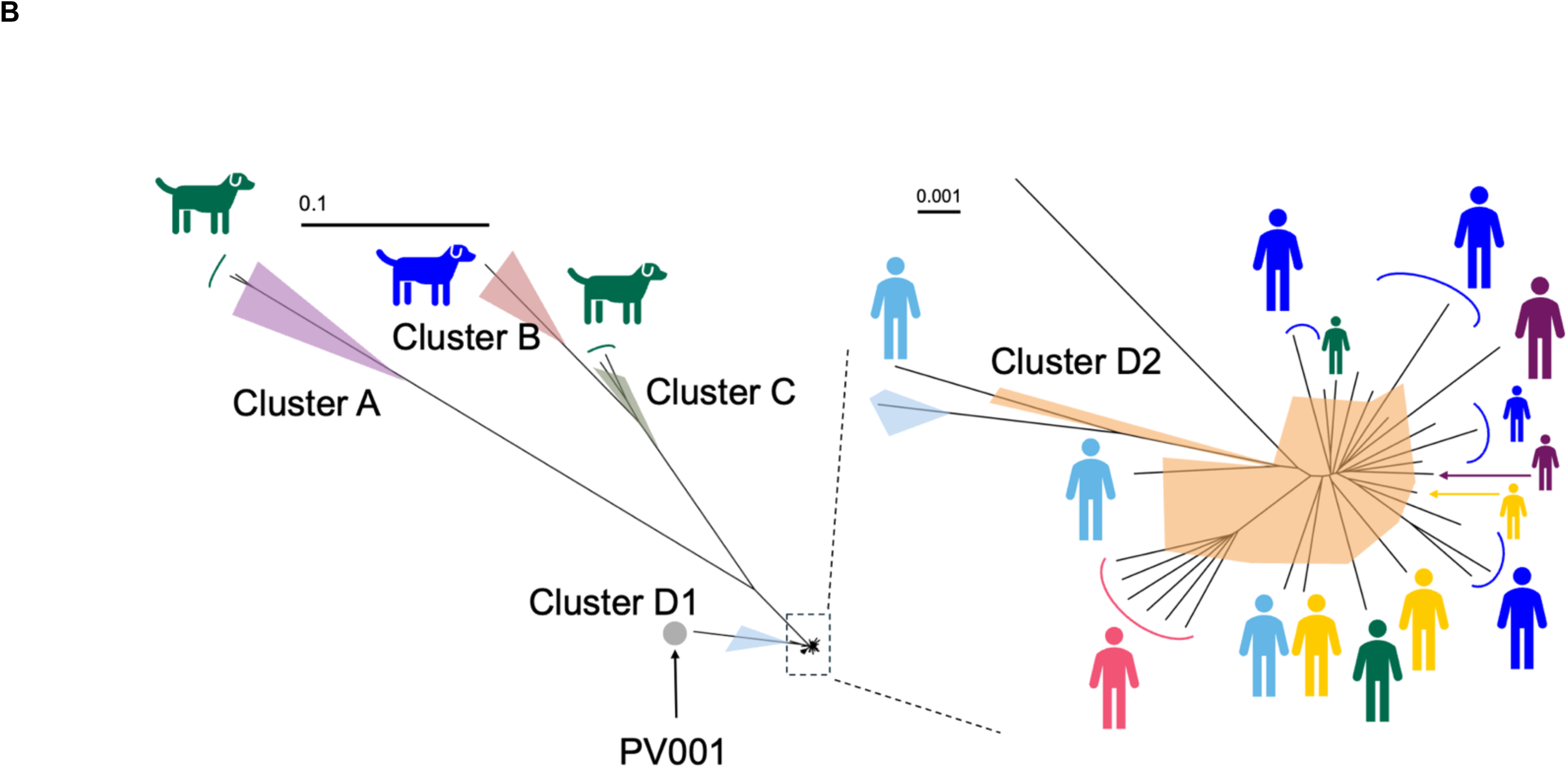
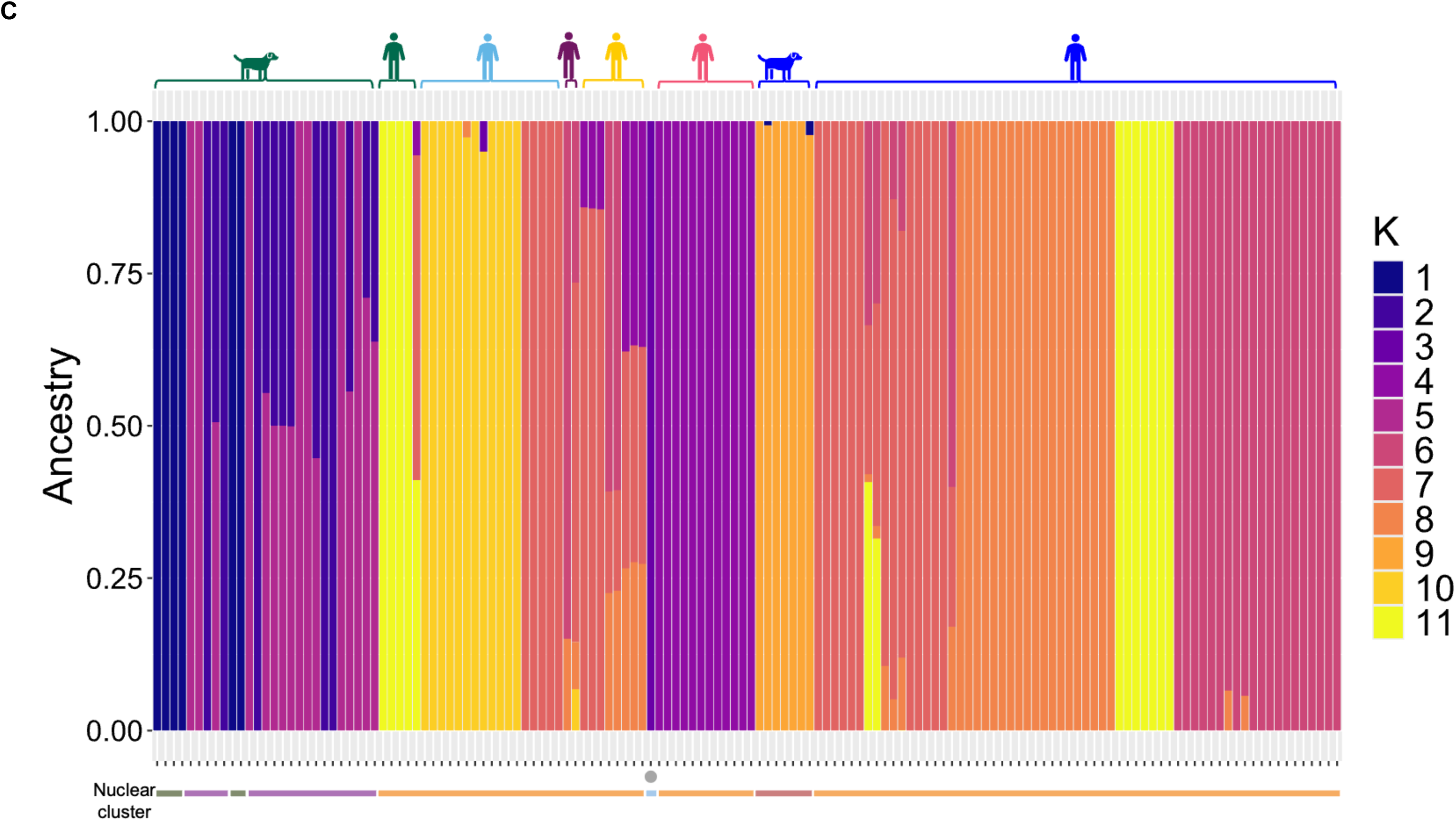
Nuclear neighbour joining (NJ) tree structure and admixture with sample reduction. (A) removal of the 15 combined samples leaving 144 iL3 and (B) also removing all but one iL3 from each host, leaving 34 iL3 samples. Comparison of these with the full tree (Figure 2B) shows that the overall structure of the trees, and the existence of the clusters A, B, C, D1 (all from dogs) and D2 (from people), is maintained. In A and B the scales are 1 substitution per 10 bases, and per 1,000 bases. (C) Admixture after removal of the 15 combined samples leaving 144 iL3s, with ADMIXTURE best supporting k = 11 groups, where 32 % (11 of 34) dog-derived samples have evidence of admixture compared to 19 % (21 of 110) for the human-derived samples. Host species and geographical location of iL3s shown at the top of the plot, and the nuclear clusters of each at the bottom. The grey dot in (A), (B) and (C) indicates the *S. stercoralis* reference PV001.

**Figure S7.**
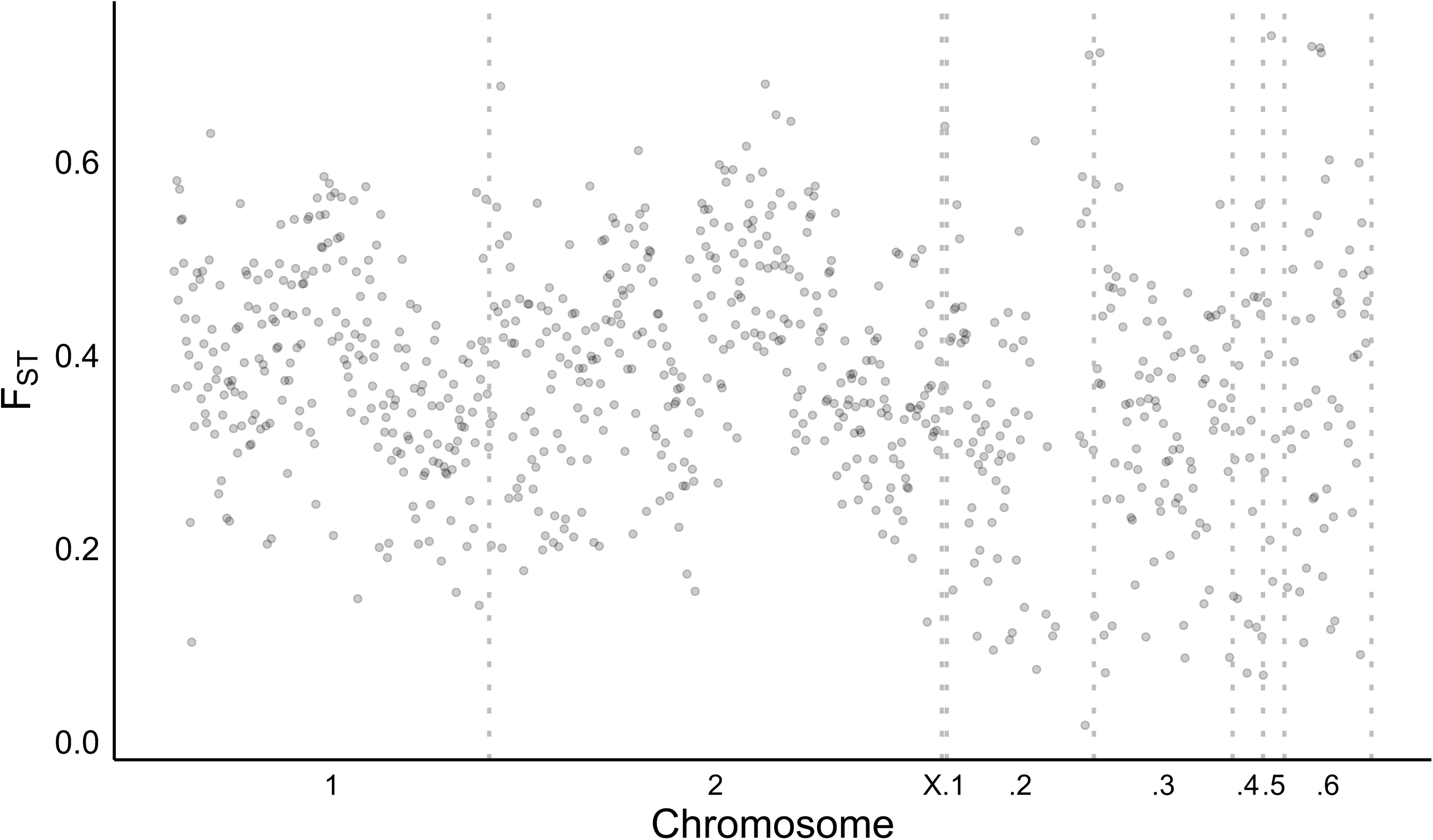
F_ST_ values across the genome. F_ST_ values in 50 kb sliding windows across the genome among iL3s from the two different host species, humans and dogs. The x-axis shows the different chromosomes of the *S. stercoralis* genome, with 6 scaffolds of the X chromosome. Grey dashed lines show the boundaries between chromosomes and scaffolds.

**Figure S8.**
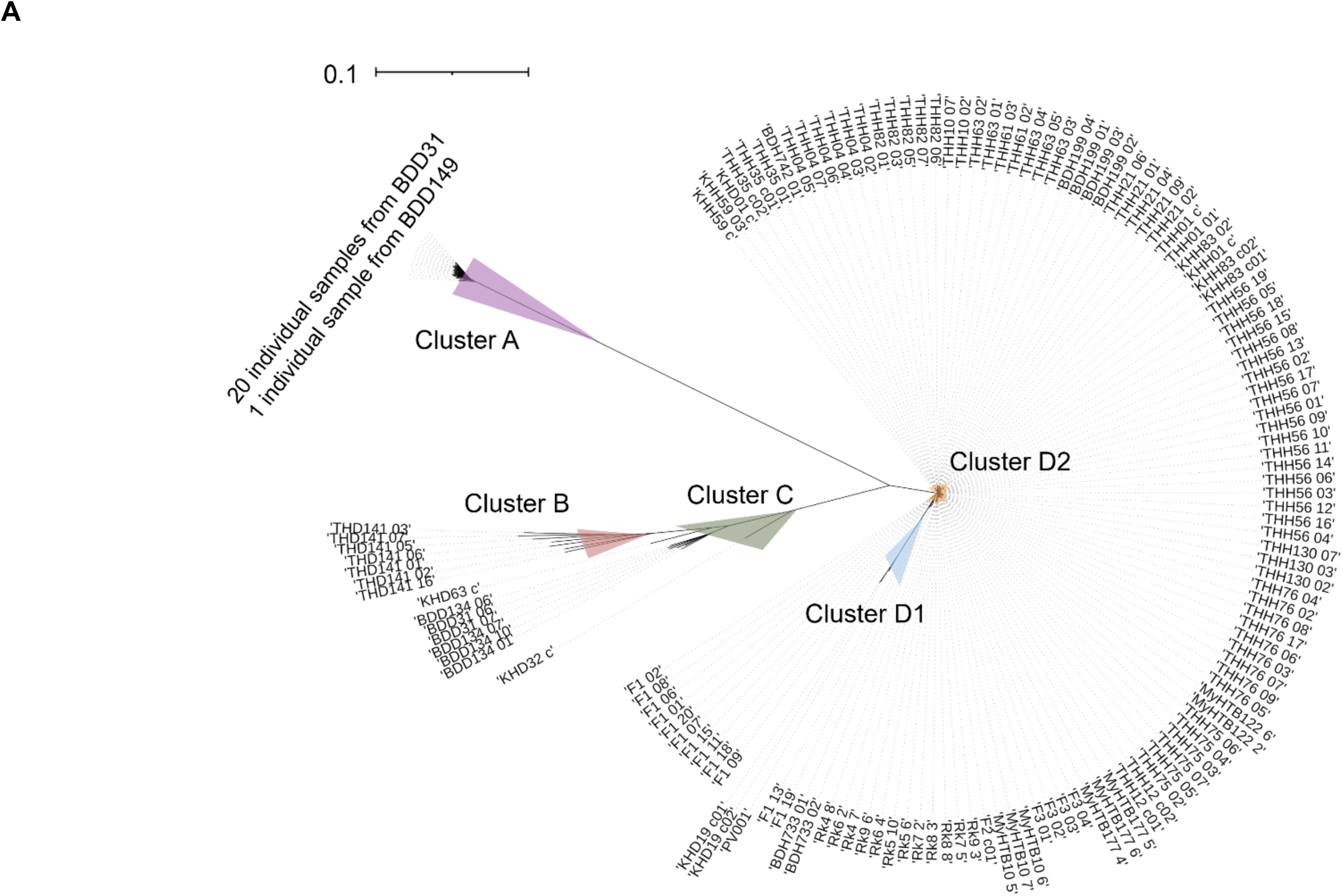
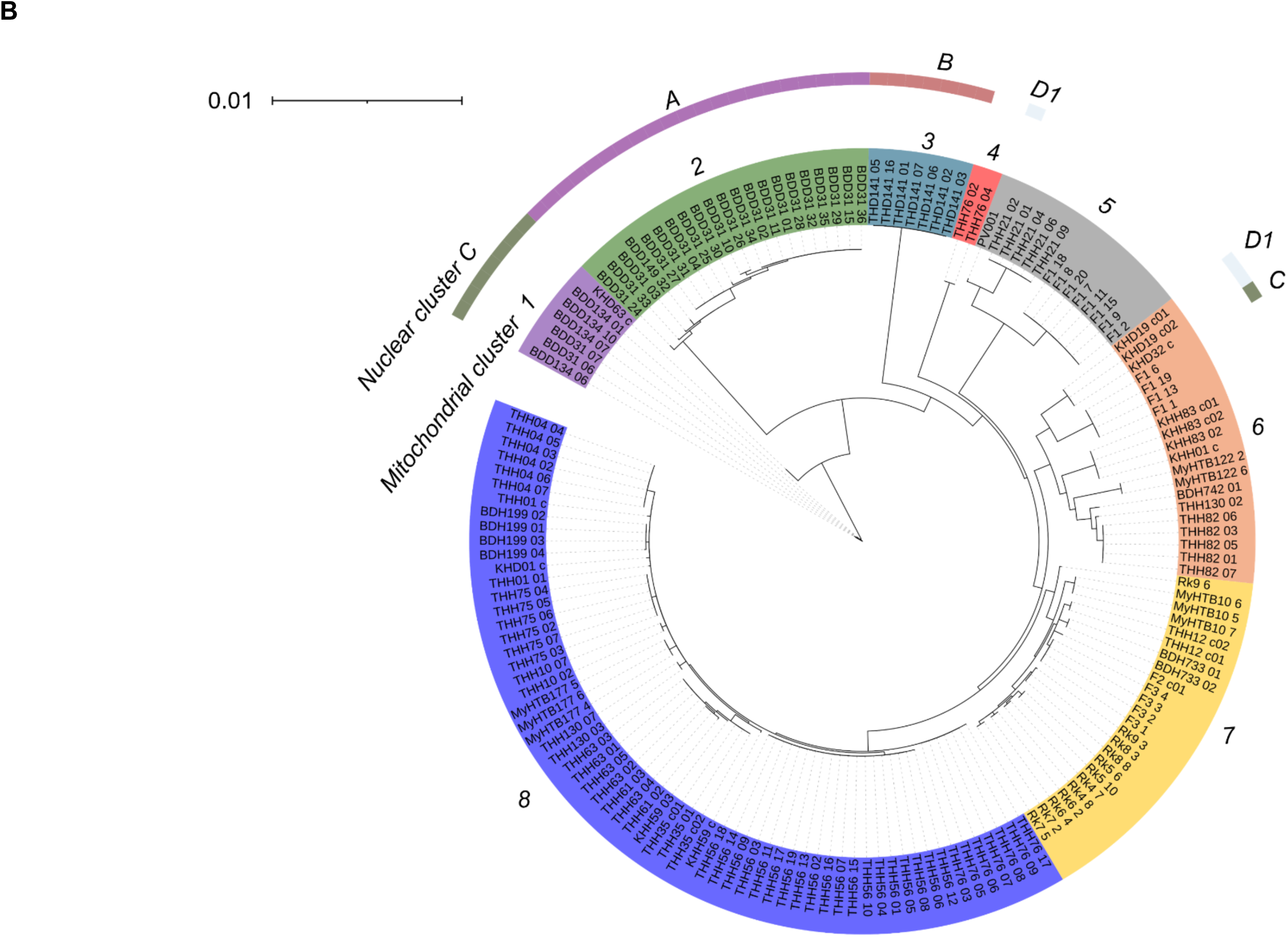
Trees with individual sample identifiers. (A) A neighbour joining tree of human and dog-derived iL3s, as main text Figure 2B, but showing the individual iL3 identifiers. (B) A maximum likelihood tree of human and dog-derived iL3s, as main text Figure 3A, but showing the individual iL3 identifiers. Scales are 1 substitution per 10 bases in (A), and 1 substitution per 100 bases in (B).

**Table S1.**
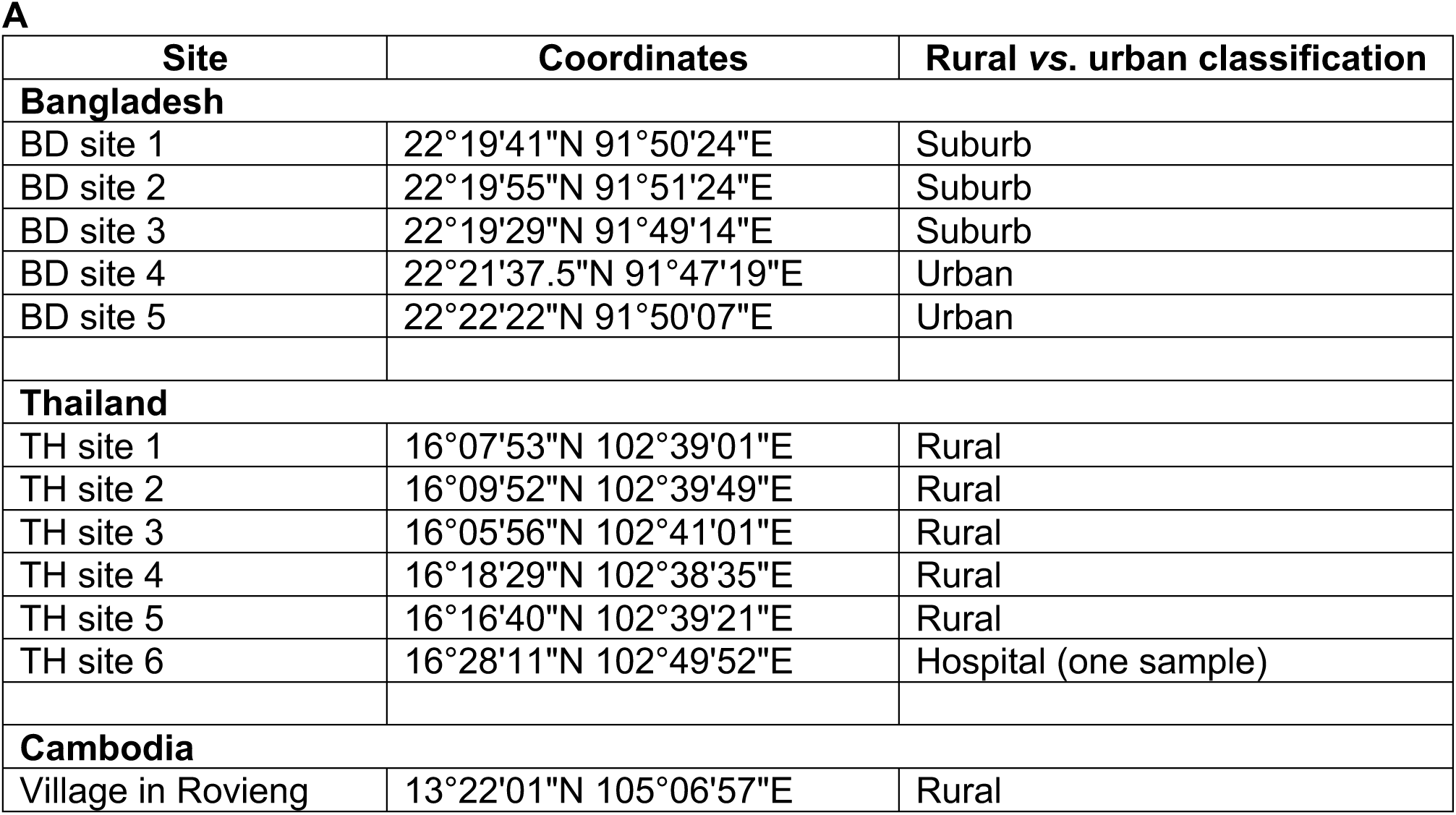
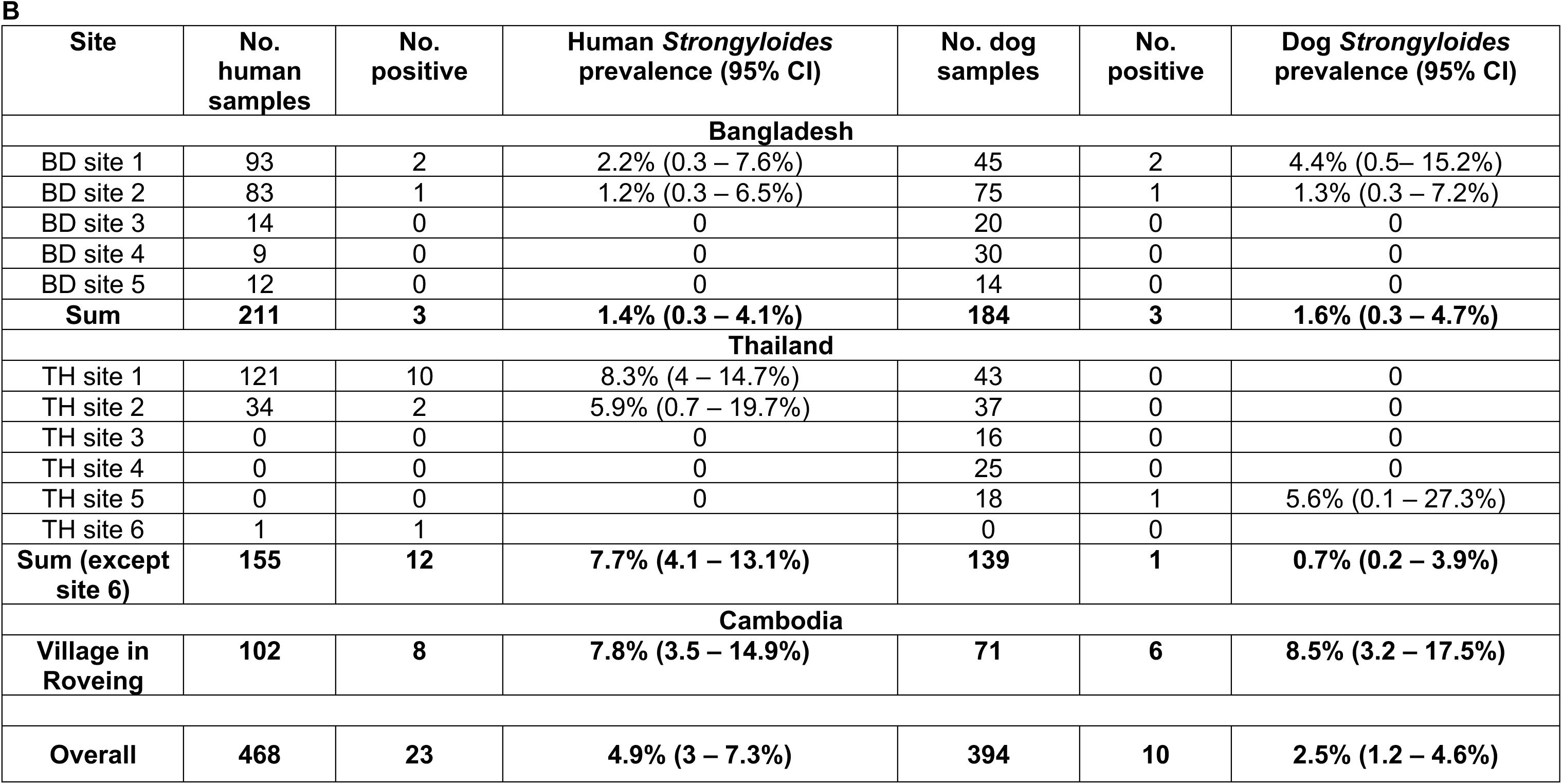
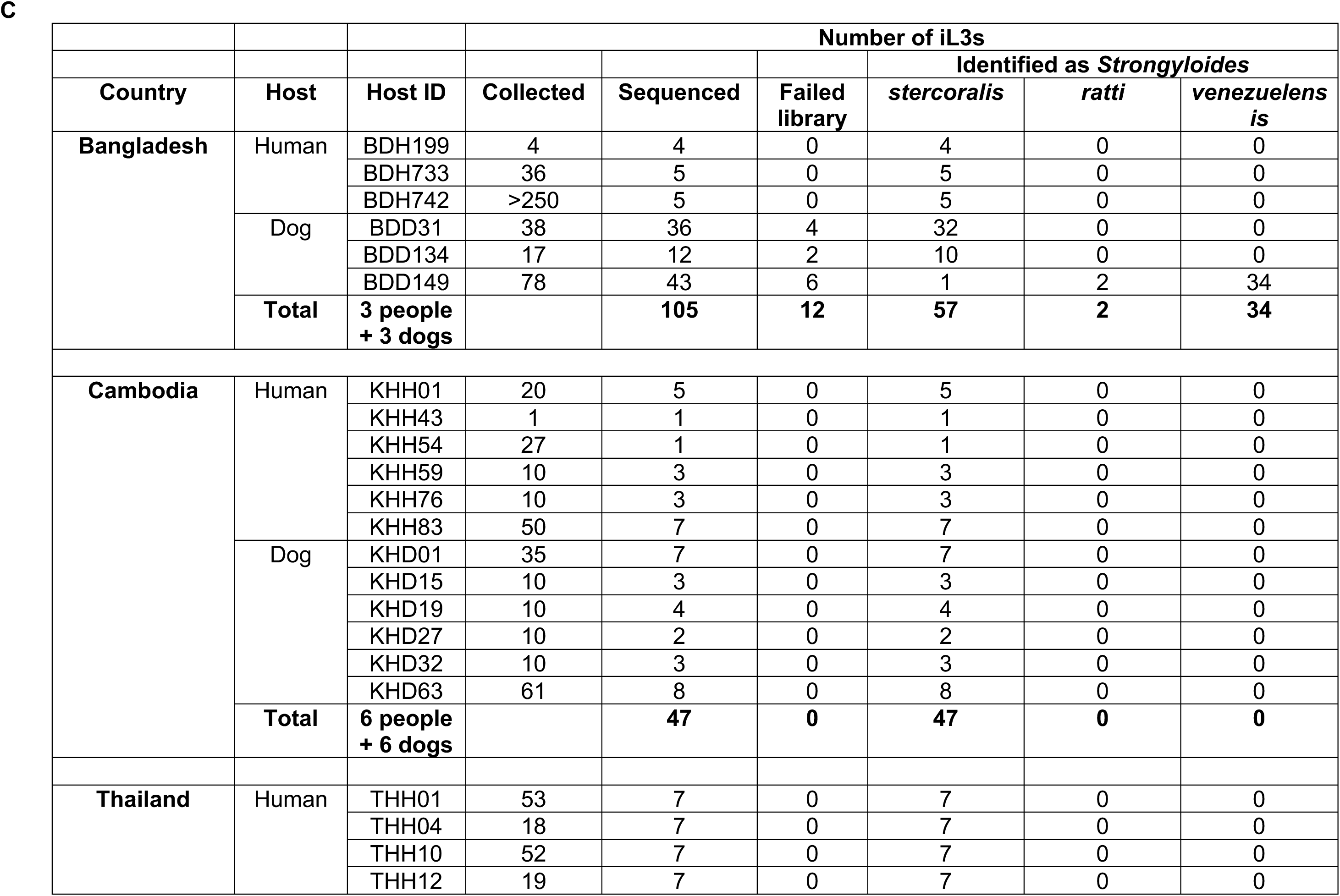
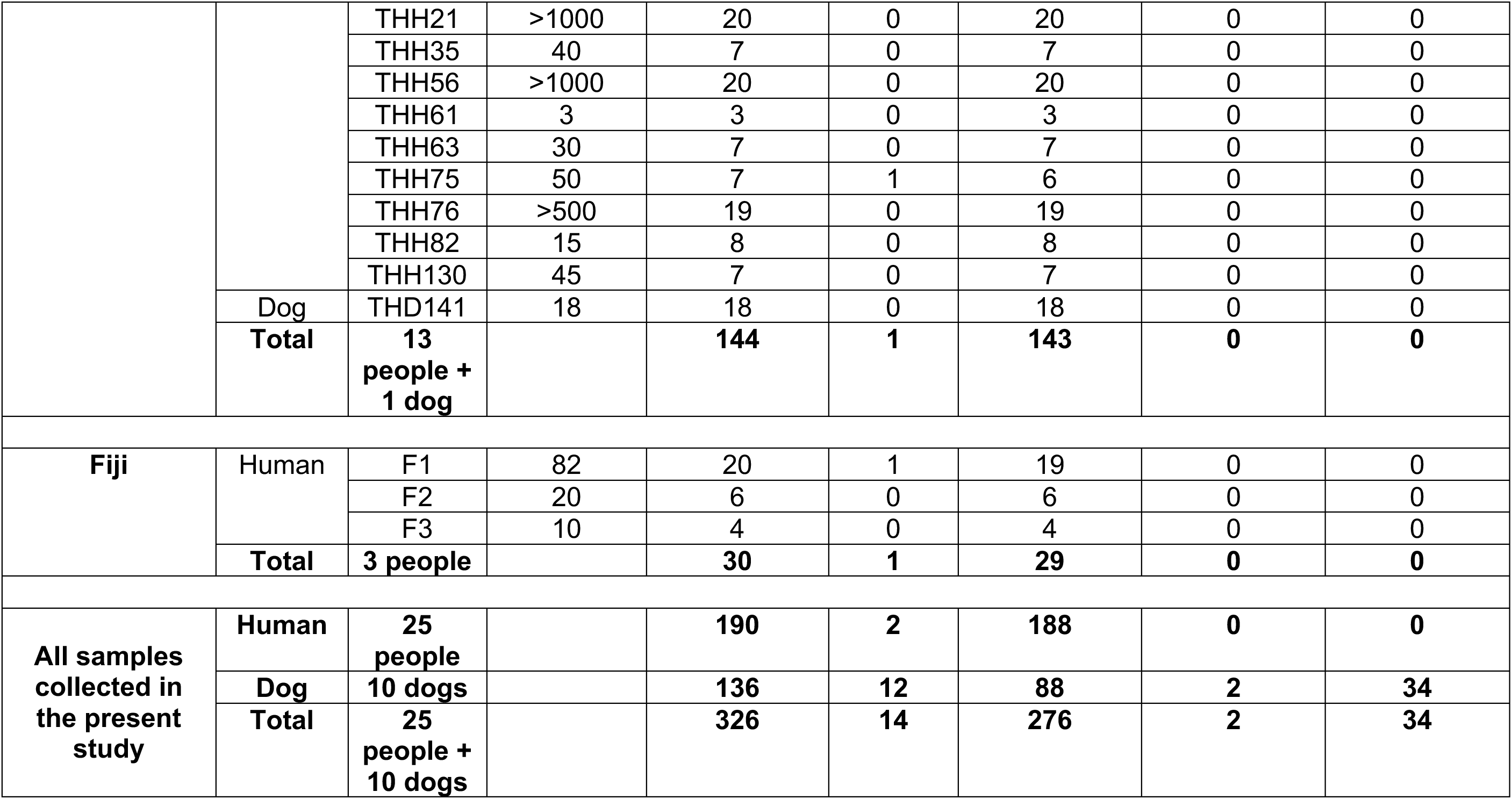
Origins of samples. (A) Location of samples sites in Bangladesh, Thailand and Cambodia. (B) Number of human and dog faecal samples, number of *Strongyloides* positives, and *Strongyloides* prevalence. (C) The number of iL3s collected and successfully sequenced from individual hosts. Failed library is where no *Strongyloides* sequence data were generated; Identified as *Strongyloides* is the Kraken-defined *Strongyloides* species identity.

**Table S2.** Combined sequence samples. The average depth and genome coverage of 56 iL3s, and then combined (shown in bold). The combined samples are named with the suffix “c” with the contributing individual host IDs in parentheses.

| Sample ID | Average depth | Genome coverage |
| --- | --- | --- |
| KHH01 01 | 4 | 41% |
| KHH01 02 | 7 | 55% |
| KHH01 03 | 11 | 68% |
| KHH01 04 | 9 | 59% |
| KHH01 05 | 25 | 70% |
| **KHH01_c(03-05)** | **52** | **97%** |
| KHH59 01 | 11 | 72% |
| KHH59 02 | 10 | 63% |
| **KHH59_c(01-02)** | **21** | **88%** |
| KHH83 01 | 14 | 77% |
| KHH83 03 | 108 | 76% |
| KHH83 04 | 44 | 63% |
| KHH83 05 | 54 | 67% |
| KHH83 06 | 3 | 45% |
| KHH83 07 | 4 | 65% |
| **KHH83_c01 (01and03)** | **123** | **94%** |
| **KHH83_c02 (04and05)** | **98** | **87%** |
| KHD01 01 | 18 | 32% |
| KHD01 02 | 26 | 34% |
| KHD01 03 | 45 | 53% |
| KHD01 04 | 8 | 20% |
| KHD01 05 | 16 | 41% |
| KHD01 06 | 9 | 24% |
| KHD01 07 | 10 | 24% |
| **KHD01_c(01,02,03,05,06,07)** | **114** | **89%** |
| KHD19 01 | 33 | 59% |
| KHD19 02 | 15 | 47% |
| KHD19 03 | 29 | 65% |
| KHD19 04 | 38 | 60% |
| **KHD19_c01 (02and03)** | **44** | **81%** |
| **KHD19_c02 (01and04)** | **70** | **83%** |
| KHD32 01 | 31 | 51% |
| KHD32 02 | 40 | 62% |
| KHD32 03 | 19 | 41% |
| **KHD32(c(01-03)** | **90** | **88%** |
| KHD63 01 | 22 | 32% |
| KHD63 02 | 21 | 26% |
| KHD63 03 | 30 | 44% |
| KHD63 04 | 16 | 30% |
| KHD63 05 | 12 | 32% |
| KHD63 06 | 24 | 36% |
| KHD63 07 | 21 | 28% |
| KHD63 08 | 8 | 15% |
| **KHD63_c(1,2,3,4,6,7,8)** | **153** | **89%** |
| F2 01 | 7 | 92% |
| F2 02 | 3 | 79% |
| F2 03 | 4 | 81% |
| F2 04 | 8 | 93% |
| F2 05 | 5 | 89% |
| F2 06 | 17 | 99% |
| **F2_c01(1,4,6)** | **31** | **99%** |
| THH01 02 | 16 | 99% |
| THH01 03 | 12 | 98% |
| THH01 04 | 13 | 99% |
| THH01 05 | 13 | 99% |
| **THH01_c(2-5)** | **53** | **99%** |
| THH12 01 | 12 | 99% |
| THH12 02 | 15 | 99% |
| THH12 03 | 10 | 98% |
| THH12 06 | 12 | 99% |
| THH12 07 | 14 | 99% |
| **THH12_c01 (1,2)** | **26** | **99%** |
| **THH12_c02 (6,7)** | **25** | **99%** |
| THH35 01 | 21 | 99% |
| THH35 02 | 13 | 99% |
| THH35 03 | 12 | 99% |
| THH35 04 | 11 | 99% |
| THH35 05 | 11 | 99% |
| THH35 06 | 11 | 99% |
| **THH35_c01 (2,6)** | **24** | **99%** |
| **THH35_c02 (3,4,5)** | **34** | **99%** |

**Table S3.** Details of sequenced iL3s that passed our QC and that were included in our bioinformatic analyses. Host ID and number of iL3s from each host from Bangladesh, Cambodia, Thailand (this study) and for Fiji, Myanmar, Japan and the *S. stercoralis* reference PV001 (all previously published data).

| Country | Host ID | Number of iL3s included in analyses |
| --- | --- | --- |
| **Bangladesh** | BDH199 | 4 iL3s |
|  | BDH733 | 2 iL3s |
|  | BDH742 | 1 iL3 |
|  | BDD31 | 22 iL3s |
|  | BDD134 | 4 iL3s |
|  | BDD149 | 1 iL3 |
|  | **3 people** | **7 iL3s** |
|  | **3 dogs** | **27 iL3s** |
| **Cambodia** | KHH01 | 1 combined sample |
|  | KHH59 | 1 iL3 and 1 combined |
|  | KHH83 | 1 iL3 and 2 combined |
|  | KHD01 | 1 combined sample |
|  | KHD19 | 2 combined samples |
|  | KHD32 | 1 combined sample |
|  | KHD63 | 1 combined sample |
|  | **3 people** | **2 iL3s, 4 combined** |
|  | **4 dogs** | **5 combined** |
| **Thailand** | THH01 | 1 iL3sand 1 combined |
|  | THH04 | 6 iL3s |
|  | THH10 | 2 iL3s |
|  | THH12 | 2 combined samples |
|  | THH21 | 5 iL3s |
|  | THH35 | 1 iL3sand 2 combined |
|  | THH56 | 19 iL3s |
|  | THH61 | 2 iL3s |
|  | THH63 | 5 iL3s |
|  | THH75 | 6 iL3s |
|  | THH76 | 9 iL3s |
|  | THH82 | 5 iL3s |
|  | THH130 | 3 iL3s |
|  | THD141 | 7 iL3s |
|  | **13 people** | **64 iL3s, 5 combined** |
|  | **1 dog** | **7 iL3s** |
| **Fiji** | F1 | 12 iL3s |
|  | F2 | 1 combined sample |
|  | F3 | 4 iL3s |
|  | **3 people** | **16 iL3s, 1 combined** |
| **Myanmar (NCBI PRJDB5112)** | MyHTB10 | 3 iL3s |
|  | MyHTB122 | 2 iL3s |
|  | MyHTB177 | 3 iL3s |
|  | **3 people** | **8 iL3s** |
| **Japan (NCBI PRJDB5112)** | RK4 | 2 iL3s |
|  | RK5 | 2 iL3s |
|  | RK6 | 2 iL3s |
|  | RK7 | 2 iL3s |
|  | RK8 | 2 iL3s |
|  | RK9 | 2 iL3s |
|  | **6 people** | **12 iL3s** |
| **Reference (NCBI ERX044031)** | PV001 | 1 samples (pool of iL3s) |
| **Total** | **31 people** | **109 iL3s, 10 combined** |
|  | **9 dogs** | **34 iL3s, 5 combined, 1 pool of iL3s (PV001)** |
|  | **Total (excluding the reference)** | **143 iL3s and 15 combined** |

**Table S4.** F_ST_ values among iL3s from different countries and host species. Samples are shown by their country code: BD, Bangladesh; KH, Cambodia, F, Fiji; JP, Japan; MM, Myanmar, TH, Thailand, and by the host code: H, human and D, dog. Dog to dog comparisons are highlighted in orange; dog to human comparisons are highlighted in blue; human to human comparisons are highlighted in green. * shows sympatric human to dog comparisons.

|  | BDH |  |  |  |  |  |  |  |
| --- | --- | --- | --- | --- | --- | --- | --- | --- |
| BDH |  | BDD |  |  |  |  |  |  |
| BDD | 0.39* |  | KHH |  |  |  |  |  |
| KHH | 0.19 | 0.38 |  | KHD |  |  |  |  |
| KHD | 0.19 | 0.37 | 0.17* |  | FH |  |  |  |
| FH | 0.17 | 0.43 | 0.15 | 0.37 |  | JPH |  |  |
| JPH | 0.31 | 0.42 | 0.36 | 0.31 | 0.20 |  | MMH |  |
| MMH | 0.16 | 0.40 | 0.18 | 0.22 | 0.12 | 0.28 |  | THH |
| THH | 0.14 | 0.53 | 0.08 | 0.62 | 0.10 | 0.15 | 0.07 |  |
| THD | 0.57 | 0.52 | 0.56 | 0.36 | 0.63 | 0.63 | 0.58 | 0.76* |

**Table S5.**
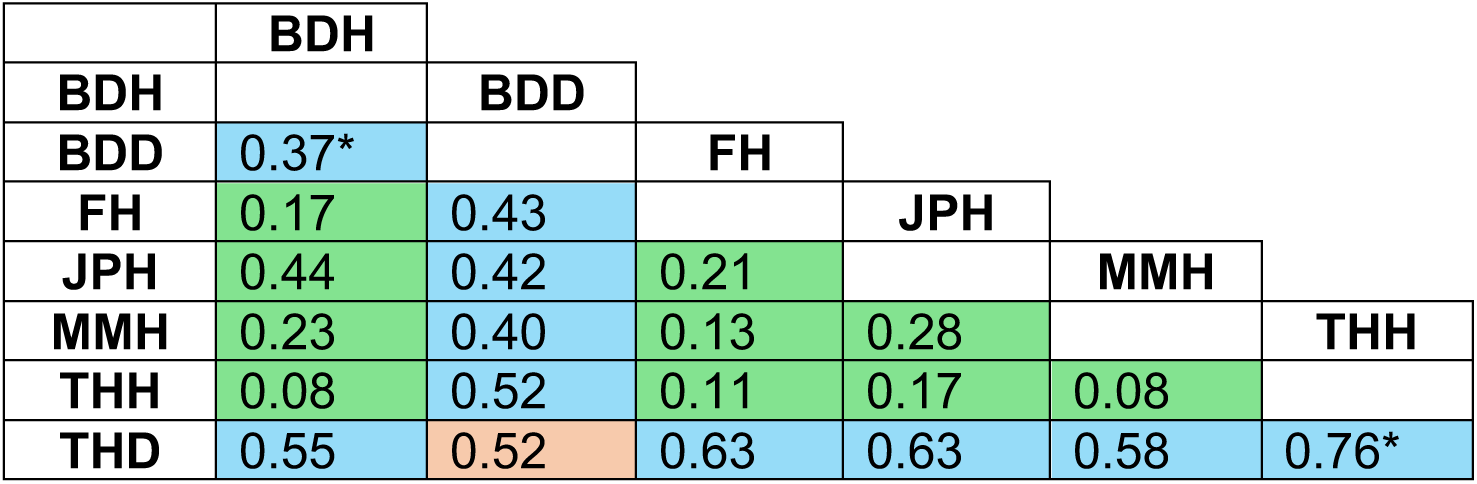
F_ST_ values among iL3s from different countries and host species with sample reduction. Samples are shown by their country code: BD, Bangladesh; F, Fiji; JP, Japan; MM, Myanmar, TH, Thailand, and by the host code: H, human and D, dog. Dog to dog comparisons are highlighted in orange; dog to human comparisons are highlighted in blue; human to human comparisons are highlighted in green. * shows sympatric human to dog comparisons.

**Table S6.** F_ST_, Admixture and dXY following modified sample inclusion. The sampled iL3s included in analyses were modified as, Modification 1 down-sampled to only a single iL3 per host, though for a single dog host (BDD31) we retained 2 iL3s that were different genotypes; Modification 2 used KING to identify identical iL3s from single hosts, which detected 35 that we then removed. Of these 35, 10 were Japanese samples and 5 Myanmar samples both of which were already published data (23, main text); Modification 3 was as Modification 2, but using dXY rather than F_ST_; Modification 4 was as Modification 1, but using dXY rather than F_ST_.

|  | **$F_{ST}$ or dXY**<br>**(median values)** | **Admixture** | **Total number of iL3s / Mean /**<br>**Maximum number of iL3s per**<br>**host** |
| --- | --- | --- | --- |
| **Modification 1** | Dog : Human = 0.65<br>Human : Human = 0.06<br>Dog : Dog = 0.17 | k = 8<br>Human only = 4<br>Dog only = 4<br>Shared = 0 | 34 / 1.1 / 2 |
| **Modification 2** | Dog : Human = 0.43<br>Human : Human = 0.14<br>Dog : Dog = 0.37 | k = 7<br>Human only = 2<br>Dog only = 4<br>Shared = 1 | 124 / 3.2 / 22 |
| **Modification 3** | Dog : Human = 0.255<br>Human : Human = 0.008<br>Dog : Dog = 0.338 | n/a | 124 / 3.2 / 22 |
| **Modification 4** | Dog : Human = 0.266<br>Human : Human = 0.009<br>Dog : Dog = 0.338 | n/a | 34 / 1.1 / 2 |

**Table S7.** Bootstrap support values. For (A) the nuclear clusters in main text Figure 2B and where values are percent of 1,000 replicates for the clades indicated, and (B) the mitochondrial clusters in main text Figure 3A, and where values are the percent of 500 bootstraps for the clades indicated.

**A**
| **Nuclear clades** | **Bootstrap support** |
| --- | --- |
| A | 100 |
| B | 94 |
| C | 85 |
| D1 | 79 |
| D2 | 82 |
| A (B ,C, D1, D2) | 100 |
| B (C, D1, D2) | 40 |
| C (D1, D2) | 98 |
| D1 (D2) | 96 |

**B**
| **Mitochondrial clades** | **Bootstrap support** |
| --- | --- |
| 1 | 100 |
| 2 | 100 |
| 3 | 100 |
| 4 | 100 |
| 5 | 100 |
| 6 | 71 |
| 7 | 65 |
| 8 | 100 |
| 1 (2,3,4,5,6,7,8) | 100 |
| 2 (3,4,5,6,7,8) | 54 |
| 3 (4,5,6,7,8) | 51 |
| 4 (5,6,7,8) | 8 |
| 5 (6,7,8) | 11 |
| 6 (7,8) | 24 |

